# Natural isolates of *Sinorhizobium meliloti* with dry, as opposed to mucoid, colony morphologies have higher soil fitness, slightly lower host fitness, and reduced nitrogen fixation

**DOI:** 10.64898/2026.10.07.757178

**Authors:** Alejandra Gil-Polo, Sohini Guha, Devan Chancey, Kayla M. Clouse, Niroshan Siva, Charles T. Anderson, Liana T. Burghardt

**Affiliations:** The Pennsylvania State University, Department of Plant Science, University Park, 16802, Pennsylvania, USA; The Pennsylvania State University, Department of Biology, University Park, 16802, Pennsylvania, USA

**Keywords:** *Sinorhizobium meliloti*, *Medicago sativa*, legume-rhizobia, mutualism, bacterial fitness, symbiosis, soil mesocosm, environmental stress, mucoid, dry

## Abstract

*Sinorhizobium meliloti* is a nitrogen-fixing soil bacterium that forms a facultative mutualism with its host alfalfa. *S. meliloti* strains exhibit diverse colony morphologies on solid culture media that span a continuum from dry to mucoid. Studies on how this trait affects symbiosis rely on mutations in exopolysaccharide II production genes; however, whether natural variation in morphology confers a selective advantage on rhizobia in their natural habitats remains unknown. Here, we used 90 mucoid and 27 dry natural isolates to test whether colony morphology associates with rhizobial fitness in the soil and in the host, or with the nitrogen-fixation benefits they provide to the plant. Colony morphologies were widely distributed across a phylogenetic tree, suggesting that morphological shifts occurred repeatedly and no single genetic variant was strongly associated with this trait. We inoculated soil mesocosms and plants with a mixture of dry and mucoid bacteria. We observed that dry rhizobia have higher fitness, particularly at 22°C and 32°C, but not under cold or drought conditions, independent of the initial rhizobial load. Conversely, mucoid bacteria have a modestly higher relative fitness in host nodules, and single inoculations showed that they tend to confer greater benefits to all three alfalfa varieties. Over time, we also observed rare, repeated conversion from mucoid to dry morphologies in soil and host plants. Taken together, our study reveals that morphological variation within a species in a single agricultural field is important for bacterial fitness in the soil and can have cascading effects on the rhizobia’s plant benefits.

## INTRODUCTION

Microbes from diverse phylogenetic groups, including *Escherichia coli* [1], *Pseudomonas aeruginosa* [2, 3], *Burkholderia cepacia* [4], *Klebsiella pneumoniae* [5], *Streptococcus pneumoniae* [6], and *Sinorhizobium meliloti* [7], can form dry and mucoid colonies on culture media. These colony morphologies are largely determined by the amount of exopolysaccharide (EPS) produced, which is governed by distinct genes and EPS types among microbes, making this one of the most striking examples of convergent evolution [8]. Authors often group colony morphology into two discrete types, dry and mucoid, which confer adaptive advantages and, therefore, higher reproductive success (fitness) in diverse habitats such as human lungs [9] or chlorinated water systems [10]. Yet both dry and mucoid morphologies often coexist and shift [1, 3], suggesting either a selective advantage to coexistence or that each morphology confers context-dependent fitness [11, 12]. However, almost all studies of the effects of colony morphology on fitness have been conducted in laboratory culture environments, limiting our understanding of how trait variation affects fitness, particularly for facultative symbionts (mutualists or pathogens) that complete their life cycles in two or more microbial habitats, alternating between free-living and host-associated stages. The selective pressures that facultative symbionts experience in both stages, and whether the trait confers fitness advantages, will ultimately drive the evolution of these microbes [13, 14]. Here, we use the rhizobia-legume model system to study how natural variation in colony morphology affects fitness both outside and inside the host.

Rhizobia are soil bacteria that form a facultative mutualism with legume plants based on nutrient exchange. They form nodules on the roots of these plants, where rhizobia fix atmospheric nitrogen, and in turn, the plant provides carbon from photosynthesis. Plants use the nitrogen for growth and development, whereas rhizobia use the carbon to reproduce and increase the number of clonal offspring derived from the (mostly) single cell that colonizes each nodule [14, 15]. Moreover, as facultative microorganisms, rhizobia can also live in the soil as free-living organisms. In natural and agricultural ecosystems, where multiple strains of the same bacterial species coexist [16, 17], competition among rhizobial strains is high [18], both in the soil and the host, and the outcomes of this competition are a major driver of rhizobial fitness. In soil, where rhizobia thrive as saprophytes or remain dormant, rhizobial strains compete for limited resources under oligotrophic conditions and in the face of various abiotic and biotic stressors [19, 20]. Furthermore, to form nodules and maximize their fitness within them, rhizobial strains must compete across multiple environments, including soil, the rhizosphere, root surfaces, infection threads, and within and across nodules [21]. Overall, in both habitats—the soil and the host plant—rhizobia will be subject to different selective pressures; therefore, traits that confer higher fitness to microbes will have a selective advantage in either habitat [21, 22]. Intriguingly, variation in colony morphology has been repeatedly observed across diverse rhizobial genera upon isolation from the field, suggesting that both morphologies coexist in nature [23, 24]. However, to our knowledge, no systematic assessment of whether this trait influences fitness in soils and hosts has ever been conducted.

To study how colony morphology influences rhizobial fitness in soil and in the plant host, we used *Sinorhizobium meliloti,* a model organism that forms an association with *Medicago sativa* (alfalfa) and has long been used as a genetic model [25]. In *S. meliloti*, mucosity is associated with exopolysaccharide production [7, 26, 27]. *S. meliloti* produces a high-molecular-weight (HMW) and low-molecular-weight (LMW) form of two types of exopolysaccharides, EPS-I and EPS-II, and decades of research have uncovered their structure, genetic bases [28, 29, 29–35], and the environmental factors that regulate their biosynthesis, i.e., quorum sensing [26, 36] and phosphate concentration [37, 38]. Single-inoculation experiments with laboratory-constructed mutants have shown that *S. meliloti*’s EPS are essential for both the successful establishment of symbiosis and for life outside the host, particularly in culture media. For example, in *S. meliloti* 1021, a model reference strain, EPS-I is crucial for establishing an effective mutualism with alfalfa, as mutants exhibit defective initiation and elongation of infection threads [39–41], but LMW EPS-II can restore this defective symbiosis [32, 42]. Outside the host, the LMW form of EPS-II is essential for biofilm formation [43], whereas the HMW form of EPS-II facilitates rhizobial movement on semisolid surfaces [37, 44]. Of the two types of EPS, EPS-II is associated with the watery-like colony phenotype on plates [37]. While we have extensive data from lab-constructed mutants, it remains underexplored whether colony morphology confers selective advantages within multi-strain populations of natural isolates.

Here, we assessed whether colony morphology, specifically dry or mucoid, was associated with rhizobial fitness in the soil, inside the plant host nodules, and whether morphology could affect the mutualistic benefit to alfalfa. We worked with a collection of 27 dry and 90 mucoid isolates from nodules of alfalfa plants grown in an alfalfa variety trial in Pennsylvania [16]. We started by quantifying EPS production for each strain and assessing their patterns on a phylogenetic tree. To assess fitness outside the host, we inoculated a mixture of 27 dry and 27 mucoid isolates into soil subjected to three temperature treatments (4°C, 22°C, and 32°C) and drought, and further broke down rhizobial fitness in the soil into two components: survival and growth (Fig. 1B). To assess fitness inside the host, we inoculated three alfalfa varieties (Vernal, Oneida, and SW4107) with mixed communities of the 117 strains and measured the rhizobial relative fitness of each strain inside the plant nodules. Finally, to measure the benefit to the host, we performed single-inoculation experiments across the three alfalfa varieties. The questions that we address are: 1) How do phylogenetic diversity and morphology relate within the *S. meliloti* strain collection? 2) Does morphology affect rhizobial fitness in the soil under varying environmental conditions? 3) Does morphology affect rhizobial fitness inside the nodules and/or provide any benefits to the plant host? We find that dry rhizobia have higher fitness in soil; their fitness in root nodules is not affected by the host, and they are less beneficial to the plant. We also observed mucoid-to-dry shifts in both habitats, and dry bacteria seem to arise independently across the phylogeny.

**Figure 1:**
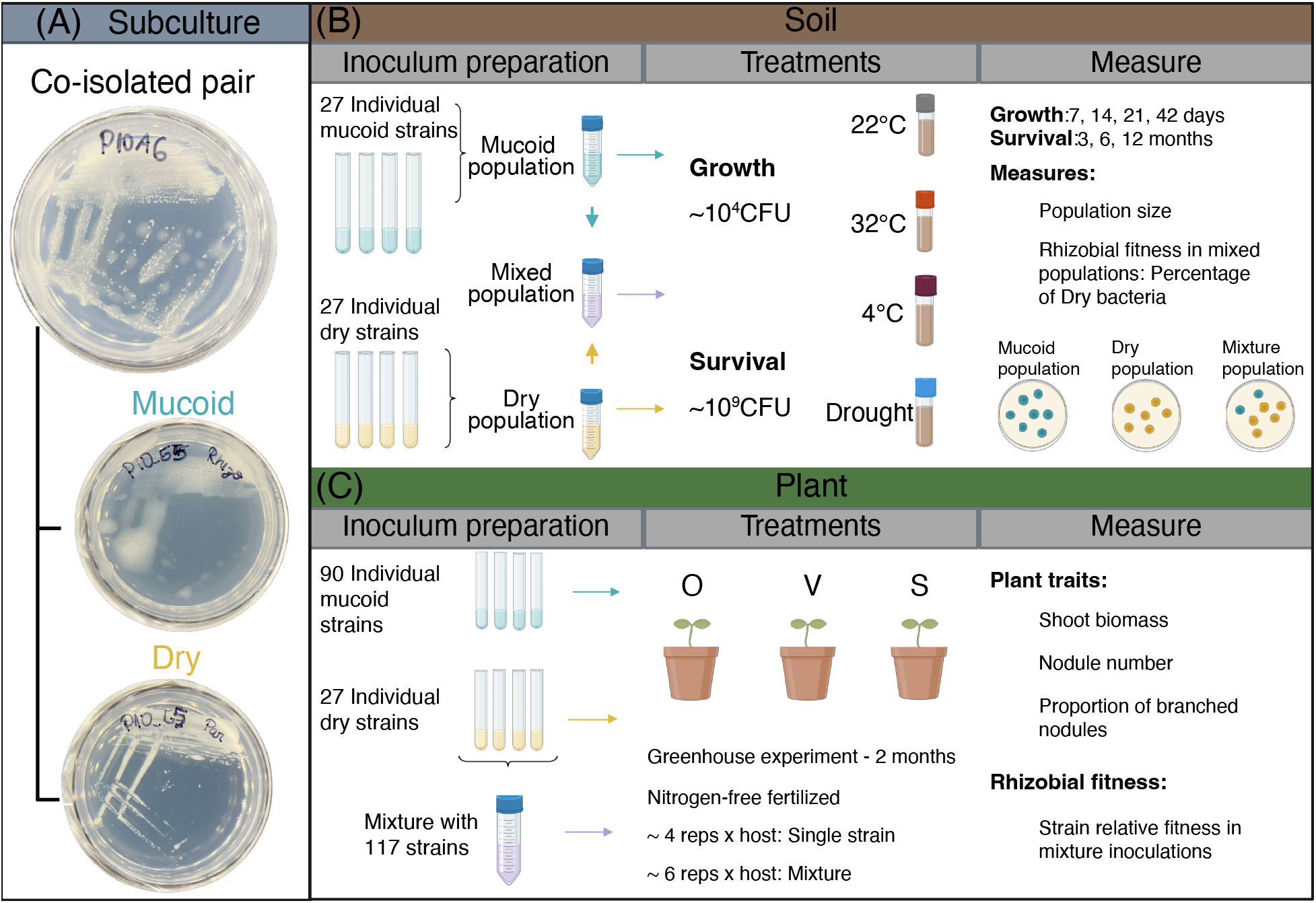
Experimental design overview. (A) Dry and mucoid bacteria were subcultured together from the GALT prospector collection and further purified. The strain with an opaque, drier morphology was classified as “Dry,” and the watery-like morphology was classified as “Mucoid.” (B) Soil selection: To prepare the inoculum, we grew 27 mucoid and 27 dry strains individually in TY. We then combined the mucoid strains to create the “Mucoid population” and the dry strains to create the “Dry population.” Both populations were mixed to form the “Mixed population.” We inoculated the soils with either a low-density inoculum (∼10^4^CFU/tube) to assess “microbial growth” or a high-density inoculum (∼10^9^ CFU/tube) to assess “microbial survival,” and exposed the soil mesocosms to 22°C (room temperature), 32°C, 4°C, and drought. Depending on the experiment (“growth” or “survival”), the harvest time points differed: shorter time points for growth (7, 14, 21, and 42 days) and longer time points for survival (3, 6, and 12 months). We later harvested the soil mesocosms, performed dilution plating, and counted colony-forming units on YEMA plates. (C) Host selection: On the plant side, we performed single-inoculation experiments with 117 strains: 90 mucoid and 27 dry. We inoculated three alfalfa varieties, Oneida (O), Vernal (V), and SW4107 (S), and kept them in the greenhouse for 2 months, fertilized with nitrogen-free fertilizer. We measured aboveground biomass, the number of nodules, and the proportion of branched nodules. We also performed a multi-strain inoculation experiment, combining all strains and inoculating the same alfalfa varieties. After harvesting the plant nodules, we followed the Select and Resequence procedure, determined the strain frequencies, and calculated relative strain fitness. Individual icons provided by Biorender.com. Image compiled and modified using Adobe Illustrator.

## MATERIALS AND METHODS

A comprehensive overview of the experiments and key analysis is provided below. Extended methodological details can be found in the supplementary materials.

### Bacterial strains and growth conditions

We isolated 117 strains from nodules of three alfalfa varieties, SW4107 (S), Oneida (O), and Vernal (V), using the General Automation Lab Technologies (GALT) Prospector (Isolation Bio, Inc.). GALT sent us the microbial isolates in 96-well plates, with each well containing a single bacterial isolate [16], putatively. During the first subculturing on Yeast Mannitol Agar (YEMA) after growing the plates in an incubator at 28°C, we observed that 32% of GALT prospector isolates exhibited two distinct colony morphologies: “Mucoid” colonies with a water-like consistency and “Dry” colonies that appeared more opaque or drier (Fig 1A). From the 117 strains isolated for the AVT collection, 90 were mucoid, and 27 were dry (Dataset S1). This strain collection was used for all experimental work. To validate that we could isolate rhizobia with distinct colony morphologies using other isolation strategies, we isolated a second group of strains by disinfecting and individually crushing 94 field-collected nodules, then performing 3 subculturing steps and recording their morphologies.

### Anthrone assay to assess exopolysaccharide production

We followed the protocol from Jones. [45] with some modifications. Briefly, we cultured 116 *S. meliloti* strains (Dataset S1) in 2 mL of *Rhizobium*-defined medium (RDM) for 8 days at 28°C and 160 RPM, with three replicates per strain. Strain 067 was not phenotyped, and strain 121 did not grow. For the anthrone assay, we centrifuged the culture for 45 min at 4°C at 10,000 g, diluted the supernatant with Milli-Q water in a 1:50 ratio, and combined 0.325 mL of this dilution with 0.675 mL of 0.2% anthrone reagent (wt/vol). We let the samples boil for 5 min, then cooled them on ice and measured the OD_620_. We estimated the glucose concentration in the samples using a calibration curve generated from glucose standards ranging from 0.156 to 20 μg. To calculate biomass, we resuspended the pellets after centrifugation and diluted them to obtain an OD_600_ between 0.05 and 0.4. Then we calculated normalized glucose equivalents, defined as Glucose/OD_600_.

### Mesocosm experiments to evaluate growth and survival

We sieved bulk soil collected from the Russell E. Larson Agricultural Research Center, mixed it with sand at a 1:1 (w/w) ratio to fill glass test tubes to a final volume of 10 g, and autoclaved the tubes. We grew 54 strains from the AVT collection in TY (Tryptone Yeast), 27 mucoid and 27 dry, at 28°C and 160 RPM for 4 days. Cultures were combined to generate a mucoid population, a dry population, and a mixed population. We inoculated the tubes with these three populations at two densities: a high density (10^9^ CFU/tube) to measure rhizobial survival and a low density (10^4^ CFU/tube) to measure rhizobial growth. We then applied four abiotic treatments: 4°C, 32°C, ∼22°C, and drought. Drought received a lower inoculum volume (adjusted to match the other treatments’ CFUs) and was the only treatment that did not receive additional water after inoculation. For the growth experiment, we harvested at earlier time points (0, 7, 14, 21, and 42 days), whereas for the survival experiment, we harvested at longer time points (0, 3, 6, and 12 months). To harvest, we added 10 mL of sterile DI water to each tube, vortexed for 1 min, and then diluted the sample 10-fold with saline solution (0.85% w/v aqueous NaCl). We then plated the dilutions onto YEMA plates and counted the colonies after ∼3 days of incubation at 28°C. In summary, N = 450: 2 inoculation densities; 4 abiotic treatments; 3 rhizobial populations; 5 time points for Growth and 4 for Survival; and 5 replicates.

### Multi-strain and single-strain inoculation experiments in plants

We sterilized, stratified in the dark at 4°C for two days, and planted three alfalfa varieties: SW401, Oneida, and Vernal. We also grew the 117 strains from the AVT collection (Dataset S1) in TY and diluted them to an OD_600_ of 0.1. For the single-strain inoculation experiment, we inoculated each alfalfa variety with 10^7^ CFU of each strain per pot (single-strain: N = 1404; 117 strains, 3 varieties, 4 reps). For the multi-strain inoculations, we mixed the OD_600_-adjusted cultures of all strains and subset four 1 mL aliquots for whole-genome sequencing to measure the initial frequency of each strain. We then inoculated the mixture in each alfalfa variety with 10^7^ CFU (multi-strain: N = 18; 3 varieties, 6 reps). We grew the plants in the greenhouse for 2 months, fertilized with nitrogen-free fertilizer. For the single inoculations, we measured dry shoot biomass, nodule number, and the proportion of branched nodules [46]. For the multi-strain, we collected all nodules from each plant, homogenized them, and submitted them for whole-genome sequencing following the protocol in [47]. We estimated the frequency of each strain in the inoculum and the nodules using Haplotype Analysis of Reads in Pools (HARP) [48], and estimated fitness as log_2_(Final/Initial frequency). More information is available in Guha et al. [16]

### Dual and single inoculation experiments

We performed dual inoculation experiments using four pairs of dry and mucoid bacteria to assess whether two morphologies can coexist within the same nodule. Those strains were also used in single inoculation experiments, where we processed all the nodules of each plant separately, to assess if morphological shifts, from mucoid to dry and dry to mucoid, were also taking place in plants. The plant host used was alfalfa variety “Oneida” (Dual inoculations: N = 28, 4 pairs, 7 reps; Single inoculations: N = 56, 8 strains, 6 reps)

### Statistical analysis of plant and rhizobial phenotypes

All statistical analyses were conducted in R version 4.4.2 (2024-10-31) and are briefly summarized here (full details in the supplementary materials). To account for population structure, we used the first principal components (PC1 and PC2) obtained after running a Principal component analysis on the Variant Call Format (VCF) file containing SNP-level variation from the 117 AVT strains (Table S1). For soil analysis, we used linear models with the aov() function, followed by an anova() call, to assess the effect of treatment and time on the percentage of dry bacteria in mixtures. To assess whether there were changes over time within each treatment, we conducted a post hoc Dunnett test against time point 0. For plant traits (shoot biomass, nodule number, proportion of branched nodules) and rhizobia relative fitness, we run a linear model to test the effects of colony morphology, alfalfa variety, their interaction, PC1, and PC2 on the average-per-strain of each of the traits. We followed with a Type II analysis of variance.

### Genome-Wide Association analysis (GWA)

We used Pyseer (v1.3.12) [49] to perform GWA using two types of genomic input: SNP variants and a presence-absence matrix. Both genomic data had an initial filter; for the presence-absence data, we filtered out genes present in less than 5% or more than 95% of strains, whereas for the SNP-based data, we filtered for variants with a higher quality than 20, a minor allele frequency lower than 0.05, and a maximum missingness of 20%. We accounted for population structure using the maximum-likelihood phylogenetic tree based on single-copy core genes shared among the 117 AVT strains [16]. When the phenotype was continuous (normalized glucose equivalent), we ran a linear mixed model (LMM); for the binary phenotype (dry and mucoid), we ran a generalized linear model. To account for multiple hypothesis testing, a significance threshold was determined using the Bonferroni correction, based on the number of unique variant patterns determined by Pyseer. We used i.TOL to visualize the tree [50].

## RESULTS

### Isolation of rhizobia with dry and mucoid colony morphologies from the field and confirmation of co-existence in nodules in the lab

Using two independent bacterial isolation methods, we found that *S. meliloti* isolates from alfalfa nodules exhibit dry and mucoid morphologies. First, the GALT-prospector method using nodule pools revealed that 32% of the individual wells from the 96-well plates sent by GALT contained both morphologies, whereas 68% contained only mucoid morphologies. This was unexpected— each GALT well typically contains a single isolate (see Materials and Methods). Second, crushing 94 individual nodules from an independent field sampling showed that 10% harbored dry rhizobia, and the rest harbored mucoid bacteria. In this second isolation, we did not observe individual nodules with both morphologies within the limits of sample size. Thus, to confirm that dry and mucoid morphologies could be co-isolated because they coexist in a single nodule, we crushed 280 individual nodules from plants co-inoculated with four pairs of co-isolated dry and mucoid bacteria, which means they were subcultured together by the GALT prospector, and observed that the percentage of mixed nodules in the 225 nodules we scored ranged from 8% to 26% across pairs (Table S2, Fig. S2B). These results demonstrate phenotypic diversity in isolates from agricultural ecosystems and co-occurrence in nodules in the lab.

### Confirmation of the association between morphology classes and EPS production

Quantification of exopolysaccharide production of the AVT collection showed that colony morphology was associated with EPS production. First, we observed that within an individual pair of co-isolated strains, the mucoid morphology produced more normalized glucose equivalent (Glucose/OD_600_) than the dry morphology (Morphology *P* = 0.002; Fig. S3A; Table S3A), validating the initial classification based on visual inspection of the subcultured plates (Fig 1A). Second, this finding held across 115 strains: mucoid strains produced higher normalized glucose-equivalent levels than dry strains (Morphology; *P* < 0.001; Table S3B; Fig. S3B). In sum, morphological classification based on YEMA plate phenotypes correlates with normalized glucose-equivalent values measured in liquid media.

### Dry bacteria are dispersed across the phylogeny, suggesting multiple shifts in morphology

A maximum-likelihood phylogenetic tree constructed from single-copy core genes, that were present in all 117 the strains we tested, showed that the dry bacteria are scattered across the tree (Fig. 2). We tested for phylogenetic signal in binary traits (D) and saw that the distribution of the morphology trait did not differ significantly from a random distribution across the phylogeny (Table S4, D = 0.86, *P* = 0.198) and was significantly inconsistent with the phylogenetically structured clustering expected under a Brownian motion threshold model (*P* = 0.001). These results suggest that the colony morphology trait shows little to no phylogenetic conservation, potentially due to multiple independent origins. Furthermore, when testing the extent to which population structure was associated with colony morphology, we found that neither principal component was significantly associated with this trait (PC1 *P* = 0.071; PC2 *P* = 0.363, Table S5A, Fig. S1). To identify candidate genes underlying this natural genetic variation in morphology, we conducted association analyses of gene presence/absence and SNP variation. We did not detect significant associations between any genetic variant and the binary phenotype (dry vs. mucoid), suggesting that this natural variation in morphology cannot be explained by a single predominant genetic change or even a handful of common genetic variants. Our original expectation was that co-isolated strains would be most closely related to each other, based on SNP differences between pairs, and that these differences could represent mutations on an essentially clonal background. To examine this possibility, we compared SNP differences between pairs (Fig. 2). We found that 4 pairs were most closely related to each other (less than 100 SNP differences), whereas the vast majority were not. Most strain pairs differed at hundreds of SNPs, with the most closely related pairs differing by 19 SNPs, while the most divergent differed by 26,423 SNPs (Table S6).

**Figure 2:**
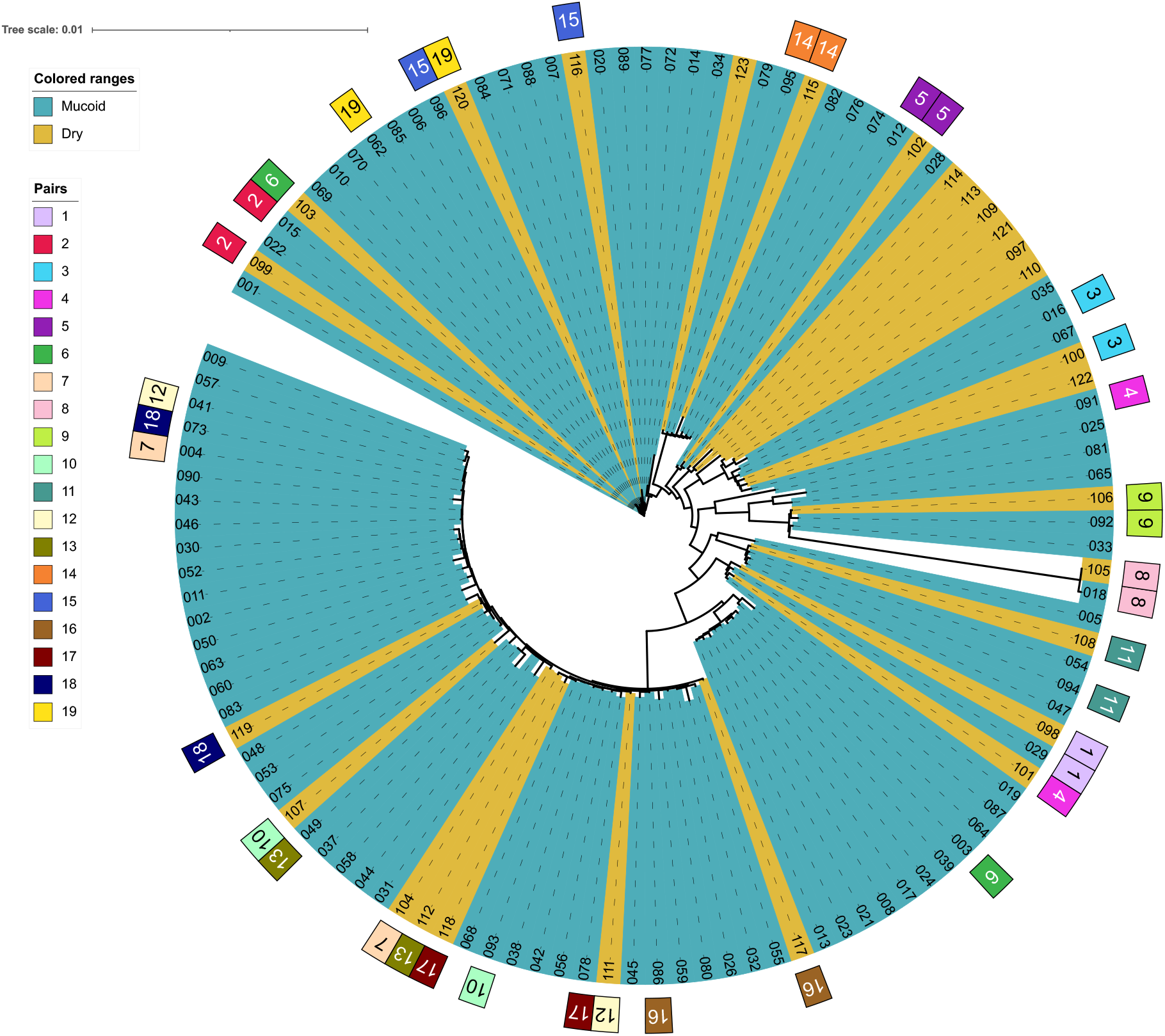
Dry and mucoid colony morphologies and co-isolated pairs are scattered across the phylogeny. A maximum-likelihood phylogenetic tree was created using IQ-TREE 3 after concatenating 3799 single-copy genes present in all 117 AVT strains. The tree shows the distribution of colony morphology, mucoid and dry, using blue and yellow colors, respectively. The tree also shows the distribution of co-isolated pairs, where squares of the same color represent a single pair (see Dataset S1). To see how many SNPs each mucoid and dry strain within a pair differed from each other, refer to Table S6. This figure was generated using iTol. See Guha et. al. (2026) for more details on the strain collection.

### In the soil, dry bacteria have higher fitness than mucoid bacteria in mixed populations

In mixed rhizobial populations grown in soil, the fitness of dry bacteria was higher in both growth and survival experiments. In the growth experiment, exponential growth occurred from day 0 to day 7, after which it plateaued (Fig 3A). However, the percentage of dry bacteria varied with time (Timepoint, *P* = 0.004, Table S7A), and significant increases in the percentage of dry bacteria were still observed after 21 days at 32°C and after 42 days at 22°C (Fig. 3C, Table S7B,C). Drought treatment did not significantly affect the fitness of dry bacteria over time (Table S7B,C), and populations kept at 4°C did not grow even after 7 months. In the survival experiments, the total population size remained constant throughout the year of the experiment (Fig. 3B), and the effect of the treatment on the percentage of dry bacteria depended on time (Treatment*Timepoint; *P* = 0.016; Table S8A). At 22°C and 32°C, the percentage of dry bacteria increased significantly from 6 months at 32°C and from 12 months at 22°C onwards (Table S8). Drought treatment, however, did not have a significant effect across time points (Table S8B), and at 4°C, the fitness of dry bacteria did not follow a trend (Fig. 3D). Taken together, these results show that the percentage of dry bacteria increases at 32°C and 22°C and after longer periods of time.

**Figure 3:**
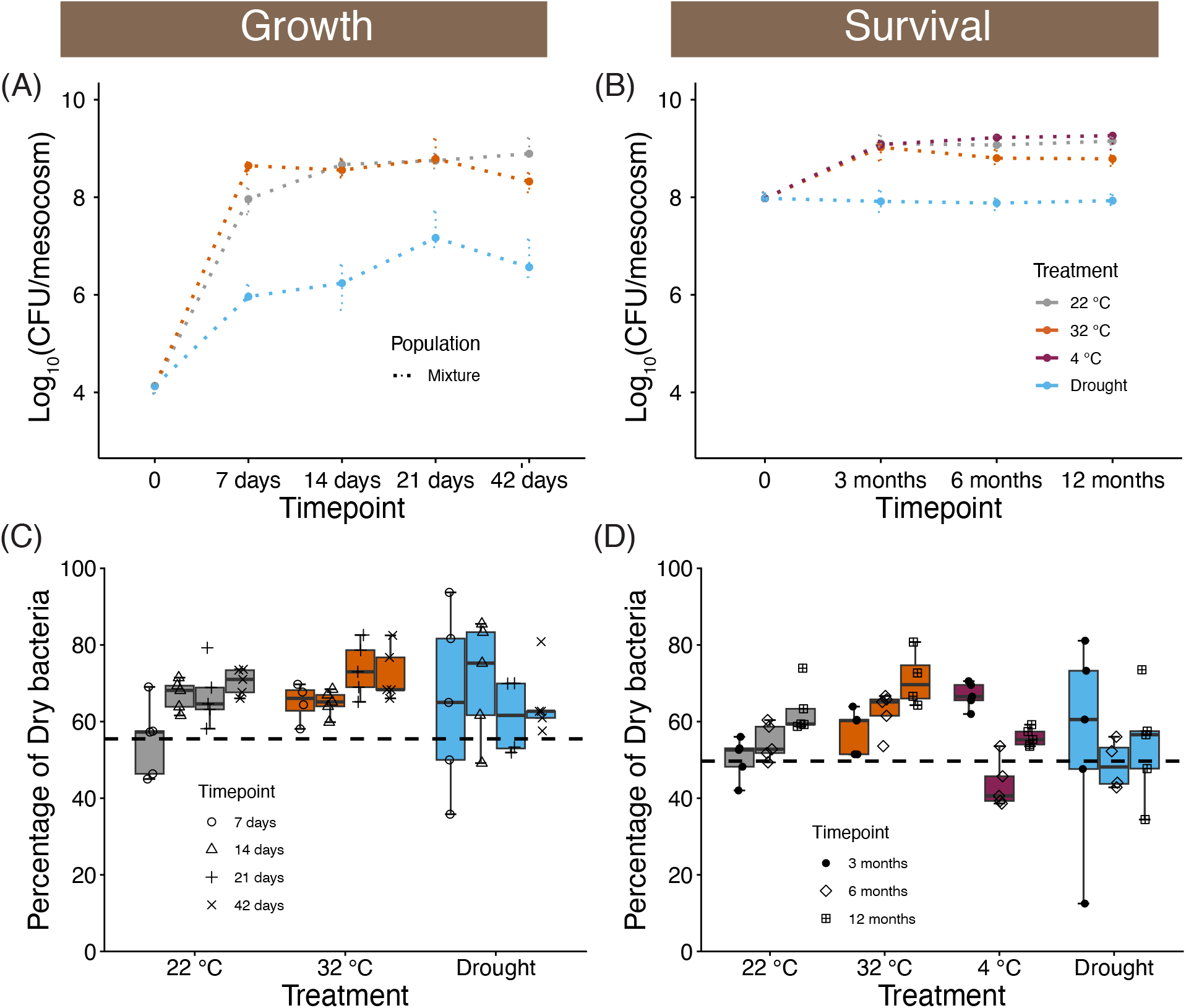
Dry rhizobia have higher fitness in the soil in comparison to mucoid bacteria in mixed populations. (A) The soil “growth” experiment with an initial low-density inoculum showed that mixtures at 22°C and 32°C exhibit similar population growth, but under drought, population size grows more slowly. At 4°C, there was no rhizobial growth. (B) The soil “survival” experiment, which started with a high-density inoculum, shows that mixed communities at 22°C, 32°C, and 4°C grow slightly at first and then maintain a steady population size, whereas under drought treatment, although there is no initial increase in number, the population size is also consistent throughout time. (C) In the growth experiment, the percentage of dry bacteria increased at later time points for treatments at 22°C and 32°C, but not for drought (Table S7). The 4°C is not represented because rhizobia did not grow under this treatment. (D) During the survival assay, the percentage of dry bacteria consistently increased at 22°C and 32°C, but not in the Drought and 4°C treatments (Table S8). Each dot in (C) and (D) represents the average of two technical replicates per rep. The black-dashed line shows the initial percentage of dry bacteria in the inoculum.

Populations of only dry or mucoid bacteria exhibited a similar pattern of population growth across most time points (Fig. S4). Interestingly, though, our calculation of population size, particularly for the mucoid community, also includes some dry bacteria that began to appear in both the growth and survival treatments (Fig. S5C and Fig. S5D). The increase in dry bacteria in soil mesocosms inoculated with mucoid bacteria is particularly high after periods longer than six months. For example, in the survival experiment at 32°C, the percentage of dry bacteria approaches 30% of the colonies after one year (Fig. S5D). These shifts from mucoid to dry, however, are very rare in the growth experiment, except under drought, where they can reach up to 15% (Fig. S5C). We did not observe transitions from dry to mucoid bacteria in the soil experiment. Taken together, our results show that mucoid bacteria can transition to a dry morphology over longer time periods, such as 6 and 12 months, particularly under warm conditions.

### With hosts, mucoid bacteria have higher relative fitness than dry bacteria and also confer additional benefits to the plants

We used a multi-strain inoculation experiment to assess rhizobia’s relative fitness in nodules of all three alfalfa varieties. After accounting for population structure (PC1 and PC2), we observed a significant effect of colony morphology (Morphology, *P* = 0.003, Table S9A). This significant effect was primarily driven by alfalfa variety SW4107; mucoid bacteria had significantly higher relative fitness than dry bacteria, whereas in alfalfa varieties Oneida and Vernal, although mucoid strains have slightly higher fitness, they are not statistically significant (Fig. 4B; SW4107, *P* = 0.015; Oneida, *P* = 0.126; Vernal, *P* = 0.086, Table S9B). When assessing the co-isolated pairs with dual inoculations, the mucoid advantage was not consistent, as we observed substantial variation in the fitness of dry bacteria among pairs—one had higher fitness than the mucoid, one had lower fitness, and two were equivalent (Fig. S6), although this experiment was limited to var. Oneida, in which mucoid and dry bacteria do not significantly differ in their fitness at the population level. Finally, sporadically, dry bacteria appeared in nodules of plants inoculated with a mucoid strain, with four out of 25 plants showing dry colonies, and these dry colonies represented a small percentage of the rhizobial population in each plant (0.05%-23.81%, Fig. S7). Our results suggest that mucoid rhizobia have a modest selective advantage within host nodules.

**Figure 4:**
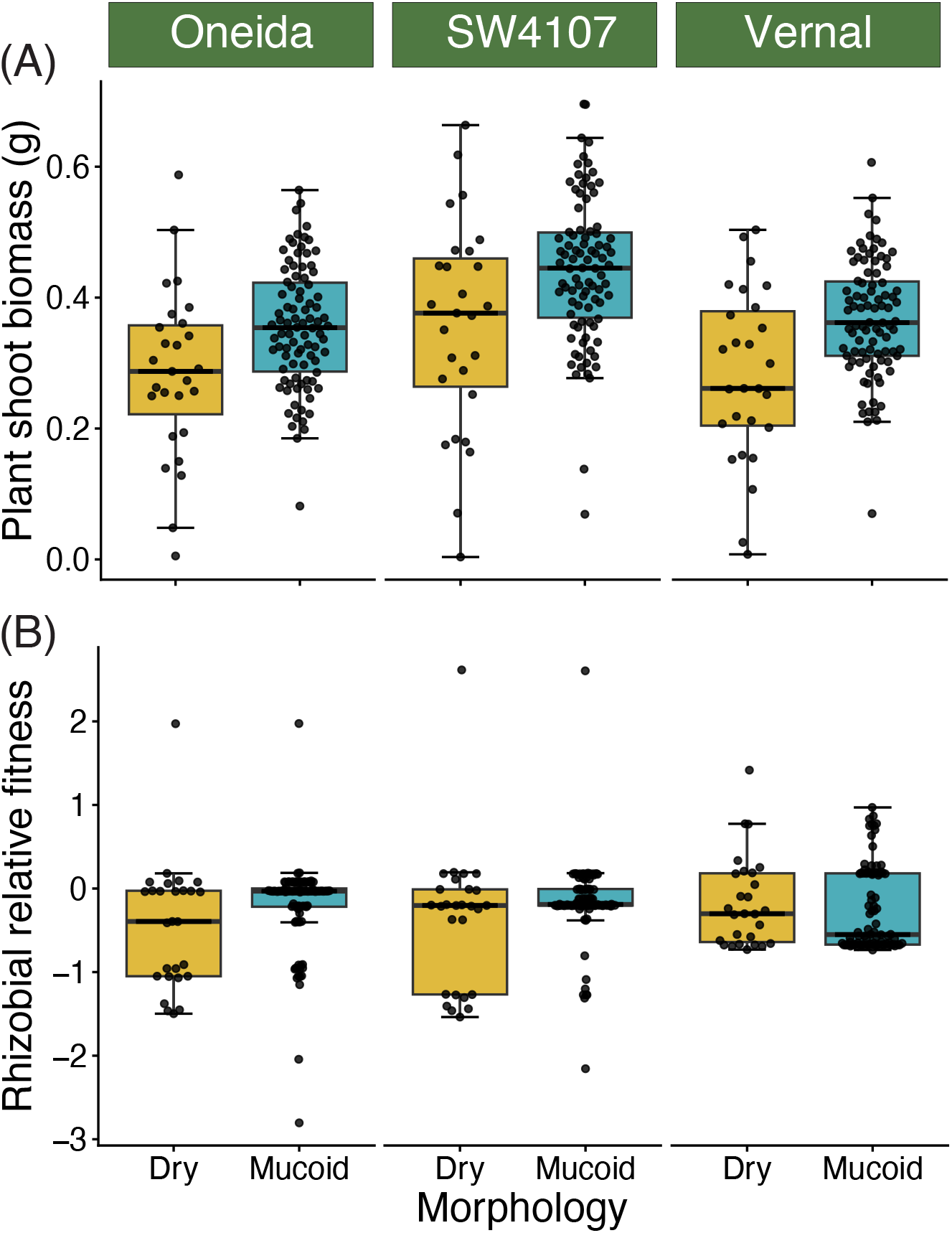
Mucoid bacteria are more beneficial and possess higher relative fitness across alfalfa varieties. (A) Single-inoculation experiments with the 117 strains show that plants inoculated with mucoid bacteria have higher shoot biomass than those inoculated with dry bacteria across the three alfalfa varieties (Table S10A). (B) Multi-strain inoculation experiments with the 117 strains show that mucoid bacteria have a statistically higher strain-relative fitness than dry bacteria inside host nodules (Table S9). Each black dot represents the average-per-strain of each trait. Yellow represents dry bacteria, and blue represents mucoid bacteria. A total of 27 dry bacteria and 90 mucoid bacteria were tested.

We used single-strain inoculation experiments to assess plant fitness. We observed that the three alfalfa varieties inoculated with mucoid bacteria had higher shoot biomass (Morphology, *P* < 0.001, Table S10A). All the strains in the collection formed nodules with the plant hosts (Fig. S8A), but the effect of the colony morphology on the nodule number depended on the host plant (Morphology*Variety; *P* = 0.043, Table S10B). We also evaluated nodule branching since previous data suggest that the proportion of branched nodules positively correlates with plant benefit [46]. However, we did not observe a significant effect of colony morphology on the proportion of branched nodules (Morphology, *P* = 0.347; Table S10C, Fig. S8B). Overall, these results indicate that mucoid bacteria are more beneficial to their hosts than dry bacteria.

## DISCUSSION

Colony morphology variation in *S. meliloti* has been observed for decades, occurring across culture media and inoculation substrates [51]. Research exploring how this trait affects symbiosis or other bacterial functions has primarily relied on lab-constructed mutants targeting EPS-II biosynthesis or strains that showed a morphological shift under *in-vitro* conditions [37, 52–55]. Here, we ask whether colony-morphology variants that occur in natural ecosystems impact bacterial fitness and mutualism benefits. We worked with an *S. meliloti* strain collection comprising 117 strains, of which 90 were mucoid and 27 were dry, isolated from alfalfa plants growing in a single agricultural field in Pennsylvania. We observed that: 1) variation in colony morphology is distributed across the phylogeny, and we could not identify a single common genetic variant that underlying this trait in our collection suggesting multiple unique variants, 2) Dry isolates have higher competitive fitness in soil at 22°C and 32°C, particularly at longer timepoints, but not in cold or drought conditions, 3) Mucoid isolates are more beneficial for the three alfalfa varieties than dry bacteria and although mucoid bacteria have higher relative fitness across all the varieties, this effect is strongest in the commercial variety SW-4107. Overall, we find that colony morphologies that increase fitness in soils tend to also decrease host benefits while having a variety-dependent effect on strain fitness in nodules.

Our findings indicate that dry bacteria arise repeatedly across diverse genetic backgrounds in the field, since morphology lacks a phylogenetic signal, and we can recapitulate their spontaneous generation in soil and, although rare, within plant nodules. Similarly, *S. meliloti* isolated from different hosts or geographic locations have been shown to form dry colonies on culture media [51, 53, 56]. We did not observe a significant association between colony morphology and either gene presence/absence or SNPs, possibly because several types of genetic variation within the same gene or across multiple genes are independently associated with the dry phenotype; therefore, none had sufficient statistical power to be detected in our GWAS. For example, lab-constructed mutants that abolish the watery-like phenotype disrupt the genes in the EPS-II biosynthetic cluster (*wgx* genes) and the *sinI*/*ExpR* quorum-sensing system [37]. Furthermore, different types of mutations in the *expR* ORF that include either the insertion of the transposable element ISrm2011-1 [7, 27, 57], or a frameshift mutation from an 11-bp deletion, or a single nucleotide substitution [53, 57] have led to a dry morphology in commercial bioinoculants or derived lines after subculturing. We cannot rule out that some variation in this trait could be due to phenotypic plasticity, since we know from previous experimental work that within individual isogenic mucoid colonies, only 30% of the subpopulation produced EPS-II [37]. Whether some of the 70% that do not produce EPS-II display a dry phenotype after subculturing remains unexplored/unknown, but results suggest it warrants additional attention. Moreover, the generation of dry morphologies has been observed in other rhizobia genera [58–60]. All this information suggests that dry bacteria can arise repeatedly in nature via diverse mechanisms, as this confers a fitness advantage, particularly under the culture conditions used in most of these studies.

Our results show that in soil, dry bacteria have higher fitness than mucoid bacteria, whether we started with a mixture of both dry and mucoid isolates or with only mucoid bacteria, with dry bacteria eventually appearing and increasing in frequency. It has been proposed that rhizobia survive in the soil for long periods by cooperating and forming biofilms [19], where EPS-II serves as a “common good” because it is an important component of the biofilm structure [43] and could support group entropy-driven mobility and sliding [44, 61]. Dry bacteria (cheaters) could emerge to exploit these cooperative structures, as they do not incur the high-energy cost of EPS-II synthesis yet still benefit from the matrix [62]. Furthermore, work in *S. meliloti* has shown that dry morphologies could exhibit higher individual motility or swarming [53, 55, 63] and are insensitive to quorum-sensing-mediated growth at high population densities [53]. These “cheaters,” therefore, would have a head start when swarming, colonizing new spaces, and scavenging for food, and would continue to grow, ignoring quorum-sensing warnings. Dry *S. meliloti* also has a selective advantage in culture media, as this phenotype can take over populations that started as mucoid bacteria after several passages [53]. However, we did not observe such drastic or rapid changes in the frequencies of population phenotypes. Taken together, our results show that even during the stationary phase in soil, when the absolute population size is not changing, evolution is occurring and selecting for phenotypes with adaptive advantages [64].

We observed that mucoid bacteria tend to confer greater benefits than dry bacteria in single inoculations in alfalfa, but colony morphology is not consistently associated with strain relative fitness. Our results demonstrate a clear plant benefit in contrast to those reported by McInnes et al. [51] and Primo et al. [27], who did not find a difference in shoot biomass when inoculating alfalfa plants with dry or mucoid strains, potentially due to low power, as they tested only 4 and 3 isolates per type, respectively. Interestingly, we recently showed that SW4107 and Oneida, but not Vernal, can weakly enrich for more beneficial rhizobia in their nodules [16]. While colony morphology has a significant effect only on SW4107, where mucoid bacteria have significantly higher fitness, the trends across the other two hosts align with this finding: although not significant, mucoid strains show higher fitness than dry strains. Furthermore, Charoenpanich et al. [53] found that dry bacteria (due to an ExpR gene mutation) have higher nodule occupancy than mucoid bacteria; but since nodule occupancy does not necessarily reflect an undifferentiated rhizobial population, which more closely reflects fitness [46], the results of our studies are not comparable. Our results point towards the importance of examining EPS-II, which is most likely the EPS behind the mucoid phenotype, and has been scarcely studied in the context of its role in symbiosis; a lot more is known about the role of EPS-I for initial stages of host infection [65] and later stages when the nodule is mature [66]. Nevertheless, it is possible that EPS-II provides rhizobia with additional protection against reactive oxygen species during symbiosis formation [67] or even against NCR peptides [68]. The mechanisms by which mucoidy increases host benefits and/or strain fitness in nodules remain to be elucidated.

Studying the effect of rhizobia colony morphology on rhizobial fitness both outside and inside the host reveals how a trait can be selected for differently in these two habitats. In the soil, where rhizobia cooperate with other rhizobial strains, for example, by forming biofilm structures, dry rhizobia are more fit; within the host, where rhizobia have formed a mutualistic relationship with the plants and are subjected to host-imposed selection, mucoid bacteria are more fit in a host-dependent manner. Our results also suggest that, following soil adaptation, *S. meliloti* can exhibit variation in colony morphology, consistent with observations in *Bradyrhizobium*, which, despite having different types of EPS, also shows high morphological variation in colonies after soil adaptation [69]. Our soil mesocosms were limited to static environmental conditions, sterile soil, and high population size. Further studies should incorporate additional complexities considered important, such as rhizobial predators [57] and the low population densities typically found in bulk soil [26, 36]. Climate change is expected to raise soil temperatures, making it important to understand how these shifts will reshape rhizobial evolution [70] and, in turn, the mutualism that legume crops depend on. Our results show that warmer soil conditions select for dry colony morphotypes, a phenotype associated with reduced plant growth. Because this symbiosis is crucial for nitrogen fixation, temperature-driven shifts towards less beneficial rhizobia could constrain plant productivity, highlighting the need to understand microbial evolutionary responses to climate change for future sustainable agriculture.

## Acknowledgments

We thank Elizabeth Paillan, Ellen Bingham, Patrick Sydow, Jennifer Harris, Amanda Jason, Andy Swartley, Gwendolyn M. Fry, Cody L. DePew, and Vyankatesh Zambare for their support during planting, plant maintenance, and harvesting in either the growth chamber or greenhouse experiments, and also during the sampling of nodules in the field. Scott DiLoreto, greenhouse manager, provided great assistance during greenhouse work. The Genomics Core Facility of the Huck Institutes of Life Sciences at The Pennsylvania State University (RRID:SCR_023645) performed whole-genome sequencing using PacBio Revio and Sanger sequencing with the Applied Biosystems 3730XL. Penn State Institute for Computational and Data Sciences (ICDS) Roar Core Facility (RRID: SCR_026424) provided computational support. This work was supported by USDA NIFA grant no. 147309 to LTB, as well as start-up funds from the Pennsylvania State University, the USDA National Institute of Food and Agriculture, and Hatch Appropriations under Project #PEN04760 and Accession #1025611. This work was also supported by NIH grant [NIH-<u>R35GM162124</u>] to LTB. This work reflects the conclusions and perspective of the authors and not necessarily those of the funding agencies.

## Author Contributions

AGP and LTB conceptualized the idea and designed the experiments. LTB obtained the funding. AGP primarily led and executed the experiments listed in this work. AGP, LTB, and SG contributed during the plant care and harvesting in both plant experiments. NS and CTA supported the exopolysaccharide measurements, and DC supported the construction of the strain collection and soil experiments. AGP, SG, and KMC analyzed the genomic data. AGP ran the statistical tests for the phenotypic data. AGP drafted the manuscript with guidance from LTB, and all coauthors reviewed and edited the manuscript.

## Data Availability

The raw sequence reads for the *Sinorhizobium* isolates are available in the NCBI Sequence Read Archive under BioProject PRJNA1482181 (Submission ID: SUB16286289). Further details on the analysis of isolate genomes are available in the preprint (Guha et al. 2026), which is currently under review. All raw data and analysis code underpinning this manuscript are in a private GitHub repository and will be revised and uploaded to a permanent repository and assigned a DOI after review and prior to publication (e.g., Ag Commons/Dryad).

