## Supplementary Material for "Natural isolates of *Sinorhizobium meliloti* with dry, as opposed to mucoid, colony morphologies have higher soil fitness, slightly lower host fitness, and reduced nitrogen fixation"

**Extended Materials and Methods:**

**Bacterial strains and growth conditions**

We assessed the colony morphology of 117 strains from our collection. Those rhizobial strains were isolated from root nodules of three alfalfa (*Medicago sativa*) varieties (SW4107, Oneida, and Vernal) using the General Automation Lab Technologies (GALT) Prospector (Isolation Bio, Inc.). The alfalfa plants had been growing at the Russell E. Larson Agricultural Research Center (Rock Springs, PA) since 2016 and were harvested in the summer of 2020. GALT sorts and captures individual cells into nanoscale microwells. They are subsequently transferred to 96-well plates with Yeast Mannitol Broth (YMB) or Reasoner's 2A Broth (R2A), from which we subcultured 371 individual wells, each putatively harboring descendants of a single strain, and purified the strains twice on Yeast Mannitol Agar (YEMA). More information can be found in Guha et al., (2026)

In the initial subculturing, we observed the coexistence of two distinct colony morphologies in a subset of plates (32%) (Fig. 1A). We assigned a morphological category (morphology) based on a qualitative comparison within each pair: the watery strain was classified as "Mucoid," while the more opaque or drier-looking strain was classified as "Dry" (Fig 1A). From the 117 strains isolated for the AVT collection (Dataset S1), 90 were mucoid, and 27 were dry. We observed that the morphological categories held up across media when we evaluated morphological consistency in other culture media, including Rhizobium-defined media (RDM) and Tryptone-Yeast (TY).

A second group of strains was isolated from root nodules using a different approach to validate whether we could isolate rhizobia with dry, mucoid colonies using other isolation strategies. In 2024, we collected 94 nodules from alfalfa varieties Oneida, Vernal, and SW4107 grown in a plot planted in 2022. We bleached each nodule individually with 4% sodium hypochlorite for 90 s and rinsed it 3 times with sterile DI water. Then we crushed each nodule with a plastic pestle and streaked it onto YEMA plates. We subcultured each strain three times, recorded its morphology, and then stored each stock at -80 °C.

For both of the microbial isolations we performed, we used Yeast Mannitol Agar (YEMA, per 1 L: 10 g Mannitol, 0.5 g $K_2HPO_4$, 0.2 g $MgSO_4.7H_2O$, 0.1 g NaCl, 0.5 g Yeast extract, 15 g agar;

adjust pH to 6.8), and for broth, we used Tryptone–Yeast extract media (TY) (per 1 L: 6 g tryptone, 3 g Yeast Extract, 0.38 g $CaCl_2$).

**Anthrone assay to assess exopolysaccharide production.**

To quantitatively assess total EPS production, we followed the protocol from Jones, (2012) with some modifications. We cultured 116 the *S. meliloti* strains (Dataset S1) in 2 mL of *Rhizobium* defined media or RDM [10 g mannitol, 100 mL RDM A stock (6 g KNO3, 0.765 g $CaCl_2$, 2.5 g $MgSO_4$.7H2O, 0.1 g $FeCl_3.6H_2O$, 1 L DI $H_2O$), 100 mL RDM B stock (10 g $K_2HPO_4$, 10g $KH_2PO_4$, 1L DI water), 4 mL of biotin stock (0.25 mg/mL), 1 mL thiamine stock (10mg/mL), 1 L DI water] for 8 days at 28 °C and 160 RPM, with three replicates per strain. We centrifuged 1 mL of the culture for 45 min at 4 °C at 10,000 g, transferred the supernatants to 2 mL Eppendorf tubes, and stored them at -80 °C. For the anthrone assay, we diluted the supernatant to 1:50 with Milli-Q water, then mixed 0.325 mL of the diluted supernatant with 0.675 mL of 0.2% anthrone reagent (wt/vol) in 98% sulfuric acid. The samples were then boiled for 5 minutes and immediately moved to ice. We then measured $OD_{620}$ by taking 200 uL of each sample and placing them in a 96-well plate. We created a calibration curve using glucose standards ranging from 0.156 to 20 µg to determine the glucose concentration in each sample. The blank and glucose calibration standards were prepared with RDM diluted to 1:50.

To normalize the production of EPS by the biomass of each strain at the time of harvesting, we estimated the biomass using the $OD_{600}$ of each pellet. We resuspended the pellets in 1.5 mL of saline solution (0.85% w/v aqueous NaCl), further diluted the suspension 1:3 in saline, and measured the $OD_{600}$ using a spectrophotometer. The purpose of the dilutions is to place most $OD_{600}$ values within the range 0.05 to 0.4, where we have observed a linear correlation between $OD_{600}$ and population size (Data not shown). We calculated Glucose/$OD_{600}$ to normalize glucose concentration relative to biomass (diluted pellet) and referred to this as normalized glucose equivalents.

**Phylogenetic tree and genome-wide association analysis**

The 117 strains in the AVT collection, as described in Guha et al., (2026), were sequenced, assembled, and annotated using the PacBio Sequel IIe platform, hifiasm v0.25.0 [3], and Prokka [4, 5], respectively. We ran the annotated genomes through Panaroo [6], which clusters genes into orthologous groups and estimates the percentage of strains carrying each cluster, thereby enabling classification of genes as core or accessory. In the panaroo analysis, the two outgroups were also included: *Sinorhizobium meliloti* 1021 and *Sinorhizobium meliloti* USDA1106. We then selected the core genes (those present in all strains) and used Panaroo's core-genome alignment of these genes as input to IQ-Tree 3 to construct the maximum-likelihood phylogenetic tree [7].

We used Pyseer (v1.3.12) to conduct the Genome-wide association (GWAS) analysis [8]. We performed associations using two types of genomic variation: gene presence/absence and single-nucleotide polymorphisms (SNPs). For presence/absence analysis, we used the presence/absence matrix generated by Panaroo as input and filtered out genes present in fewer than 5% or more than 95% of the strain collection. We went from 12591 groups, defined as clusters of orthologous genes, to 3760. For the SNP variation analysis, we used the variant calling format (VCF) file generated by mapping the whole-genome sequences of our strain collection to the in-house reference genome AVT118 using bwa mem v0.7.17 [9]. We used vcftools v0.1.17  [10] to filter for variants with variant quality greater than 20, minor allele frequency greater than 0.05, and missing in at most 20% of samples. After this filtering step, we kept 47393 out of 136,566 sites. To enable rapid regional querying of the VCF file, we used bgzip (htslib, v1.24) [11] to compress the file and tabix (htslib, v1.24)[12] to generate spatial indexes. To account for population structure, we used the script phylogeny_distance.py, part of the Pyseer scripts, to convert the maximum-likelihood phylogenetic tree that only included the 117 AVT strains (made with 3799 single-copy core genes, not strain outgroups included in this tree) into both a similarity-based matrix (-lmm) and a distance-based matrix (without -lmm). For the SNP variation analysis, PySeer skipped the multiallelic sites.

For the GWA analysis, we used two types of phenotypic data: binary data, in which the phenotype was either mucoid or dry, and continuous data, in which the phenotype was the normalized glucose equivalent. To assess the genetic associations with the binary phenotype (Dry and Mucoid), we ran a generalized linear model with the distance-based matrix as a fixed effect to account for population structure. To assess the genetic associations with the continuous phenotype (normalized glucose equivalent), we fitted a linear mixed model with the similarity-based matrix as a random effect. We used PySeer pattern hashing to count unique patterns of identical genetic-variant distributions across samples. Later, we used the Pyseer script count_patterns.py to determine the number of patterns and, therefore, compute a Bonferroni-corrected significant p-value threshold.

To assess whether colony morphology shows phylogenetic signal, meaning whether related strains tend to share similar traits because of a common ancestor, consistent with a single ancestral origin versus multiple independent origins, we calculated the D statistic [13] using the phylo.d function in the R package caper, with 1000 permutations against both a random-association null (D= 1) and a brownian-motion clumped null (D=0). To run this test, we started with the maximum-likelihood phylogenetic tree constructed from the 2977 single-copy core genes of the 117 AVT strains, plus three outgroups often used as reference strains (*S. medicae* USDA1037, *S. meliloti* Rm1021, *S. meliloti* USDA1106). We then dropped *S. medicae* USDA1037 and used the root() function from the ape package [14] to root the tree with the remaining two reference strains. After rooting the tree, we resolved any polytomies using the multi2di() function.

**Soil mesocosm experiments to evaluate growth and survival.**

We collected bulk soil from the Larson Agricultural Research Center and allowed it to air-dry at room temperature. To remove small pieces of roots or stone, we sieved the soil. Then we filled 25 × 100 mm glass test tubes with 10 g of 1:1 sand and soil (w/w) and autoclaved them once using a liquid cycle at 121 °C for 30 min. We applied three temperature treatments: cold (4 °C, in the fridge), warm (32 °C, in the growth chamber), and room temperature (~22 °C, in the lab), as well as a drought treatment (~22 °C, in the lab). We weighed the tubes biweekly, and when they had lost 1.5 g, we added 1.5 ml of DI water, except in the drought treatment, where no additional water was added throughout the experiment.

We inoculated soil tubes from each treatment with three distinct rhizobial populations. One made up of 27 randomly selected mucoid strains (mucoid population), another made up of 27 dry strains (dry population), and a mixture of the 54 dry and mucoid strains (mixed population) (Fig. 1B, Dataset S1). We used two different inoculum densities, a low-density inoculum (~ $10^4$ CFU/tube) and a high-density inoculum (~ $10^9$ CFU/tube), to assess rhizobial reproductive success during growth and survival, respectively. In both cases, we grew each strain in TY for 4 days in a shaker at 28 °C and 160 RPM. For growth experiments, we pooled mucoid and dry strains separately by combining 500 µL of each strain. Both suspensions were vortexed, centrifuged at 5,000 x g for 15 min at 6 °C, washed twice with saline solution, and resuspended to reach an $OD_{600}$ of 0.1. To create the mixed population, we combined the dry and mucoid suspension. We estimated that 10 µL of a 0.1 $OD_{600}$ suspension corresponds to ~$10^6$ CFU, so we diluted the suspensions to prepare an inoculum containing $10^4$ CFU per soil tube. For the survival experiment, we pooled 2.2 mL of each strain's culture to generate the dry and mucoid populations separately, and 1.1 mL of all strains' culture to generate the mixed population. We estimate that each culture had ~$10^9$ CFU/mL. Each population was centrifuged and washed as described above, then resuspended in saline to prepare an inoculum containing $10^9$ CFU per soil tube. We added 2 mL of inoculum to each tube for the 22 °C, 32 °C, and 4 °C treatments, and 0.5 mL for the drought treatment. In both growth and survival experiments, the inoculum for the drought treatment is more concentrated, so although they receive a lower initial volume, the number of cells is the same as in the other treatments.

We harvested the growth experiment at earlier time points (0, 7, 14, 21, and 42 days), and the survival experiment at later time points (0, 3, 6, and 12 months). Overall, we harvested 5 tubes per population per treatment, per timepoint. To harvest each tube, we added 10 mL of sterile DI water and vortexed each tube for 1 min. Then, we took 100 µL of the soil suspension and diluted it 10-fold in 900 µL of saline solution. We then plated a sufficient volume of the dilution to yield 20 to 80 colonies on YEMA. We counted the mucoid and dry-looking colonies.

**Multi-strain and single-strain inoculation experiments in plants**

More details of this experiment can be found in Guha et al. (2026). Briefly, we planted Vernal, SW4107, and Oneida in a 1:1 mix of sand and turface (wt/wt). For the single inoculations, we used 90 mucoid bacteria and 27 dry bacteria (Dataset S1); for the multi-strain inoculations, we combined all 117 strains. To measure the initial frequencies of each strain in the multi-strain inoculum, we subset 4 aliquots of 1 mL each, centrifuged at 18 000 × g for 2 min, removed the supernatant, and stored the pellet at -20 °C. In both single- and multi-strain cases, we inoculated each pot with ~$10^7$ CFU. We had four replicates of each alfalfa variety inoculated in single inoculations, and six replicates for the multi-strain inoculations (single-strain N = 1404; 117 strains, 3 varieties, 4 reps; multi-strain N = 18; 3 varieties, 6 reps). The plants were maintained in the greenhouse and fertilized with a nitrogen-free fertilizer, and were harvested 60 days post-inoculation. We cut the shoots, let them dry at 60 °C for a week, and then measured aboveground biomass. We also collected all nodules from each plant, counted the number of nodules per plant, and calculated the proportion of branched nodules [15]. For plants inoculated with the multi-strain inoculum, we collected all nodules from each plant and processed them as described in [16] to calculate the rhizobial relative fitness. Briefly, we homogenized each nodule pool using a tissue homogenizer, enriched for the undifferentiated rhizobia population using two centrifugation steps, and extracted DNA [17]. DNA was sequenced on an Illumina MiSeq (2×150 bp), trimmed with TrimGalore! (v0.4.1, www.bioinformatics.babraham.ac.uk/projects/trim_galore/), aligned to in-house *S. meliloti* reference strain AVT118, and frequencies were estimated using Haplotype Analysis of Reads in Pools (HARP) [18], and fitness was estimated as log2(Final/Initial frequency). The strain frequency of strain 018 on one of the Vernal sampled plants was 0, and the log adjustment resulted in –Inf, so we replaced that value with "–8".

**Dual and single inoculation experiments:**

To assess whether the co-isolated pair of dry and mucoid strains could co-exist within the same nodule in the lab, we co-inoculated alfalfa variety "Oneida" with four pairs of co-isolated strains. The pairs were: 015 and 099, 029 and 098, 016 and 100, 091 and 101. Additionally, to assess whether mucoid bacteria could shift to dry and vice versa within host nodules, we performed single-inoculation experiments. The alfalfa seeds were disinfected one day before planting with 10% commercial bleach, then rinsed, placed on sterile filter paper in petri dishes, and left to germinate overnight at room temperature. The substrate used to grow plants consisted of 70% Sunshine Mix, 20% Turface, and 10% sand. This substrate was transferred to autoclave bags and autoclaved in a liquid cycle at 121 °C for 30 min, left on the bench to cool overnight, and then autoclaved again the next day. We then filled the pots and planted the alfalfa seedlings. Fresh colonies were grown for one day in 2 mL of TY in the shaker at 28 °C and 160 RPM. Then we diluted the culture in fresh TY and adjusted the $OD_{600}$ to 0.2. We used the relationships between CFU and $OD_{600}$ observed in our calibration curves for each strain (Data not shown) to combine the diluted $OD_{600}$ with 0.85% saline solution to prepare a final inoculum containing $10^7$ CFU in 20 mL, the volume added to each pot. To create the dual inoculum, we mixed equal parts of the final inoculum from each mucoid and dry strain in each pair. We then performed a 10-fold dilution of the only-mucoid and only-dry inoculum in saline solution and plated them on YEMA to count colonies and estimate the initial population size. We had 6 reps per single inoculation per strain and 7 reps per dual inoculation per pair. The plants then grew for two months in a growth chamber without any extra nitrogen fertilization.

To assess the rhizobia population size in each plant, we processed all nodules from plants inoculated with either dry or mucoid bacteria. All nodules per plant formed an individual nodule pool, which was disinfected with 4% sodium hypochlorite for 2 min and then rinsed 3 times with DI water. The disinfected nodules were transferred to a 15 mL Falcon tube containing 3 mL of saline solution and homogenized using a tissue homogenizer [16]. Then, we used 100 µL to prepare 10-fold dilutions and plated them to count colonies and assess their colony morphologies. To evaluate whether individual nodules were occupied by dry, mucoid, or both colony morphologies, we randomly selected 10 nodules per plant from plants co-inoculated with two strains. We disinfected each nodule with 4% sodium hypochlorite for 90 s, rinsed it, crushed it with a micropestle, resuspended the resulting suspension in 0.85% saline solution, and streaked the contents onto Yeast Mannitol Agar to assess the morphology of the microbe inside each nodule.

**Statistics and Data Analysis:**

All statistical analyses were conducted in R version 4.4.2 (2024-10-31). To account for genome-wide variation among the 117 strains, we ran a Principal Component Analysis (PCA) using the VCF file generated with AVT118 as the reference. For the PCA, we used the prcomp() function, after centering the data but not scaling it, [prcomp(data, center = TRUE, scale. = FALSE)]. This VCF file contained 136,566 SNPs; after filtering for missing data, we ran PCA on 96,206 SNPs. We added PC1 and PC2 as fixed-effects predictors to account for genome-wide variation in statistical models. These two PCs explained 42.55% and 17.59% of the variation, respectively (See the variation explained by the top 10 PC in Table S1 and Figure S1). To assess the effects of genome-wide variation on the morphology (Mucoid and Dry), we fit a generalized linear model with binomial distributions using the glm() function from the stats package [glm(Morphology ~ PC1 + PC2, data, family = "binomial")] followed by Anova() from the car package [19] to perform a Type II analysis of deviance with likelihood ratio tests [Anova(model, type = "II", test.statistic = "LR")]. To assess genome-wide variation using continuous normalized glucose equivalent data as the trait, we used a linear model lm() from the stats package [lm(Normalized Glucose equivalent ~ PC1 + PC2)] followed by an analysis of variance with the anova() function [anova(model)].

To assess whether strains classified as mucoid had higher normalized glucose equivalents than dry bacteria within pairs and across the entire AVT collection, we used the per-strain average of normalized glucose equivalents for the 38 paired strains and for 115 out of the 117 strains in the AVT collection, respectively. For the comparison using only the paired strains, we used a linear mixed model with morphology as a fixed effect and pair as a random effect using the lmer() function from the lmerTest package [20], [lmer(Normalized Glucose equivalent ~ Morphology + (1|Pairs))]. The variance attributed to Pair was extracted from the model, and to test whether including Pair as a random effect significantly improved model fit relative to a model without it, we used a likelihood ratio test with the ranova() function. For the comparison across the entire AVT collection, we run a One-Way ANOVA with the lm() function, using morphology as the only predictor [aov(Normalized Glucose equivalent ~ Morphology)], followed by an analysis of variance anova().

In the analysis of the soil mesocosm experiments, we observed that the percentage of dry bacteria in the mixtures for both growth and survival follows a normal distribution, as indicated by the histograms. Therefore, to test whether the percentage of dry bacteria in the mixed community is associated with treatment (22°C, 32°C, 4°C, and Drought) and harvesting time, we ran a Two-Way ANOVA using the aov() function, [aov(Percentage of Dry bacteria ~ Treatment*Time)], followed by anova(), [anova(model)]. This two-way ANOVA was run separately for the growth and survival experiments. Then, we subset the dataset by treatment and ran a One-Way ANOVA to assess the effect of timepoint within each treatment, [aov(Percentage of Dry bacteria ~ Time)]. We then ran a post-hoc Dunnett test using the glht() function from the multcomp package in R [21], [glht(model, linfct = mcp(Timepoint = "Dunnett")] to assess whether the percentage of dry bacteria was significantly different in the later timepoints than at timepoint “0”.

For the analysis with hosts, we tested whether morphology and alfalfa variety affect the traits shoot biomass, nodule number, the proportion of branched nodules, and rhizobium relative fitness. We used the average-per-strain of each of these traits as the response, in a linear model with the lm() function, followed by an analysis of variance with the Anova(Type = “II”) function from the car package, which evaluates each effect after accounting for all the other effects in the model. Each model included morphology, alfalfa variety, and their interactions, as well as PC1 and PC2, to account for genome-wide relatedness among strains. When evaluating the rhizobia relative fitness trait, we therefore subset the analysis by Variety and then ran a linear model (lm()) followed by an analysis of variance (Anova(), Type II), using Morphology as a fixed effect and accounting for genome-wide relatedness effects.

**Table S1:** The 10 principal components (PC) of the principal component analysis (PCA) run on the variant calling format (VCF) files with the SNP-level variation of the 117 strains and the percentage of variance each one represents (Percent Variance %). These VCF files were created using the in-house AVT118 strain as the reference genome. The PCA was run to account for population structure.

| PC | Percent Variance (%) |
|---|---|
| PC1 | 42.55 |
| PC2 | 17.59 |
| PC3 | 10.39 |
| PC4 | 7.89 |
| PC5 | 4.71 |
| PC6 | 4.47 |
| PC7 | 3.31 |
| PC8 | 2.24 |
| PC9 | 1.64 |
| PC10 | 1.51 |

**Table S2:** Percentage of dry, mucoid, and mixtures (both morphologies) in the individual nodules of plants co-inoculated with four pairs of dry and mucoid strains. Strains 015, 016, 029, and 091 are mucoid, whereas strains 098, 099, 100, and 101 are dry. The percentages were calculated with respect to the number of nodules from which we extracted data, which is reported in "**Number of nodules**"

| Pair | Number of nodules | % Mucoid | % Dry | % Mixture |
|---|---|---|---|---|
| 091_101 | 58 | 68.966 | 22.414 | 8.621 |
| 015_099 | 52 | 36.538 | 46.154 | 17.308 |
| 016_100 | 65 | 49.231 | 33.846 | 16.923 |
| 029_098 | 50 | 18.000 | 56.000 | 26.000 |

**Notes:** We assessed the morphology of the microbes recovered from single nodules of plants co-inoculated with 4 pairs of dry and mucoid bacteria (Pair) and streaked in YEMA plates. **Pair**: The strains that were co-isolated from the GALT prospector. **Number of nodules:** The number of nodules from which we extracted data, because they had either mucoid, dry, or mixed morphologies. We started with 70 nodules per pair, but some did not grow or contained contaminants (displaying yellow or red growth). We are reporting only the number of nodules for which we were able to extract data, and this number was used as the denominator to calculate the percentages for Mucoid, Dry, and Mixture.**% Mucoid:** the percentage of nodules that had only the mucoid morphology. **% Dry:** the percentage of nodules that had only dry bacteria. **% Mixture:** The percentage of nodules that had Mixture (both dry and mucoid bacteria).

**Table S3:** Mucoid bacteria have higher normalized glucose equivalent than dry bacteria, when comparing co-isolated strains (A) and across the whole population (B)

(A)

| Term | Estimate | Std. Error | df | t_value | p_value |
|---|---|---|---|---|---|
| (Intercept) | 6.205 | 3.189 | 33.279 | 1.945 | **0.060** |
| MorphologyMucoid | 13.961 | 3.811 | 18.000 | 3.663 | **0.002** |
| Random effect: Pairs (variance) | 55.266 | - | - | - | - |
| Random effect: Pairs (LRT vs. no random effect) | - | - | 1 | 1.535 ($\chi^2$) | 0.215 |

**Notes**: To validate that our classifications of dry and mucoid between the co-isolated pair were associated with exopolysaccharide production, we ran a linear mixed model using the per-strain average of normalized glucose equivalent as the response variable, morphology as the fixed effect, and Pairs as the random effect. **Term**: Tells what the model is describing. "MorphologyMucoid" is the main result. "Random effect: Pairs (Variance)" row reports the variance that is attributable to Pair. "Random effect: Pairs (LRT vs. no random effect)" is the likelihood ratio test that compares the model with and without the random effect. **Estimate**: For the fixed effect, it is the effect size, and for the random effect, it is the variance. **t_value:** For the fixed effect, it is the test statistic. For LRT, the chi-square statistic ($\chi^2$) is used. Abbreviations: **Std. Error** (Standard Error), **df** (Degrees of freedom), **p_value** (P-value)

(B)

| Term | Df | Sum Sq | F value | Pr(>F) |
|---|---|---|---|---|
| Morphology | 1 | 2618.549 | 31.666 | **< 0.001** |
| Residuals | 113 | 9344.426 | NA | |

**Notes**: To validate that mucoid bacteria produced higher normalized glucose equivalent than dry bacteria across our entire collection, we calculated the per-strain average of normalized glucose equivalent and then tested Morphology by running a linear model aov() followed by an analysis of variance ANOVA, anova(). Abbreviations: **Df** (degrees of freedom), **Sum Sq** (Sum of Squares values), **Pr(>F)** (P-value)

**Table S4:** We tested for a phylogenetic signal in the morphology trait and found that it was more similar to a random pattern than to a clumped pattern.

| N_tips | D_estimate | P_random_D1 | P_brownian_D0 | Permutations |
|---|---|---|---|---|
| 117 | 0.861 | 0.198 | **0.001** | 1000 |

**Notes:** We used the phylo.d() function from the caper (comparative analysis of phylogenetic and evolution) package in R. **N_tips:** We had 117 strains in the phylogenetic tree. **D_estimate** (D) measures how a binary trait (in this case, dry and mucoid) is distributed on a tree to measure the phylogenetic signal. Since our D is near 1, the trait is distributed randomly across the tips of the tree. **P_random_D1** (P-random) is the p-value that tests whether the trait's distribution is different from a purely random association across the phylogeny. Since the p-value is greater than 0.05, the trait is distributed randomly across the tree. **P_brownian_D0** (the P-Brownian) is the p-value for testing whether the trait's distribution differs significantly from that of a Brownian motion model, which refers to a clumped phylogenetic pattern. A p-value lower than 0.05 means that we reject the Brownian motion. **Permutations**: Number of permutations of the binary trait across the tips of the phylogenetic tree.

**Table S5:** Evaluate the effect of genome-wide variation on (A) morphology, and (B) normalized glucose equivalent.

(A)

| Term | LR Chisq | Df | Pr(>Chisq) |
|---|---|---|---|
| PC1 | 3.268 | 1 | *0.071* |
| PC2 | 0.826 | 1 | 0.363 |

**Notes:** We ran a generalized linear model with a binomial error distribution and a logit link using the glm() function, followed by an analysis of deviance using likelihood-ratio tests with the Anova() function. The response variable was morphology: Mucoid and Dry, while the predictors were PC1 and PC2. **Term** (name of the exploratory variable**), LR Chisq** (Likelihood Ratio Chi-Square). **Df** (Degrees of freedom), **Pr(>Chisq)** (P-value)

(B)

| Term | Df | Sum Sq | F value | Pr(>F) |
|---|---|---|---|---|
| PC1 | 1 | 589.210 | 6.427 | **0.013** |
| PC2 | 1 | 1106.404 | 12.069 | **0.001** |
| Residuals | 112 | 10267.362 | NA | |

**Notes:** We ran a linear model with lm(), followed by an analysis of variance with anova(). The response variable was the normalized glucose equivalent. In both cases, PC1 and PC2 are the two principal components from a Principal Component Analysis (PCA) run on the SNP genetic variants. **Term** (name of the variable), **Df** (Degrees of freedom), **Sum Sq** (Sum of Squares), **P(>F)** (P-value)

**Table S6**: Number of SNP mismatches when comparing co-isolated pairs. We compared the number of mismatching sites after comparing the variant calling format (VCF) files of the 117 AVT strains in our collection. These VCF files were created after using the AVT118 strain as the reference genome.

| **Pairs** | **Mucoid strain** | **Dry strain** | **SNP differences** |
|---|---|---|---|
| 3 | 016 | 100 | 515 |
| 4 | 091 | 101 | 15858 |
| 5 | 012 | 102 | 246 |
| 6 | 003 | 103 | 26423 |
| 7 | 004 | 104 | 224 |
| 8 | 018 | 105 | 315 |
| 9 | 092 | 106 | 509 |
| 10 | 093 | 107 | 176 |
| 11 | 094 | 108 | 379 |
| 12 | 041 | 111 | 130 |
| 13 | 049 | 112 | 19 |
| 14 | 095 | 115 | 294 |
| 15 | 096 | 116 | 405 |
| 16 | 086 | 117 | 75 |
| 17 | 078 | 118 | 84 |
| 18 | 073 | 119 | 79 |
| 19 | 062 | 120 | 300 |
| 2 | 015 | 099 | 298 |
| 1 | 029 | 098 | 308 |

**Notes: Pairs** (Number assigned to each Pair that was co-isolated in the GALT prospector). **Mucoid strain** (The code of the Mucoid strain); **Dry strain** (The code of the Dry strain). **SNP differences** (number of SNP differences between strains).

**Table S7:** Statistical models to assess the effect of treatment and timepoints in the percentage of dry bacteria in the mixtures in growth experiments (soil) using (A) Two-Way ANOVA, (B) One-Way ANOVA, and a (C) Post-Hoc Dunnett test.

(A)

| Term | Df | Sum Sq | F value | Pr(>F) |
|---|---|---|---|---|
| Treatment | 2 | 164.089 | 0.850 | 0.433 |
| Timepoint | 4 | 1699.690 | 4.400 | **0.004** |
| Treatment:Timepoint | 8 | 770.438 | 0.997 | 0.449 |
| Residuals | 55 | 5311.678 | NA | |

**Notes**: Linear model followed by a Two-Way ANOVA to evaluate the effect of the Treatment and the Timepoint on the average of the percentage of dry (the average was determined among technical replicates of each biological replicate or soil tube). **Term** (predictor used and residuals), **Df** (Degrees of freedom), **Sum Sq** (sum of squares), **F value** (F stats), and **Pr(>F)** (P-value)

(B)

| Treatment | Term | Df | Sum Sq | F value | Pr(>F) |
|---|---|---|---|---|---|
| 22 °C | Timepoint | 4 | 959.624 | 5.395 | **0.004** |
| 22 °C | Residuals | 19 | 844.941 | NA | |
| 32 °C | Timepoint | 4 | 935.502 | 6.885 | **0.002** |
| 32 °C | Residuals | 18 | 611.407 | NA | |
| Drought | Timepoint | 4 | 575.002 | 0.671 | 0.620 |
| Drought | Residuals | 18 | 3855.330 | NA | |

**Notes**: One-Way ANOVA to assess the effect of the Timepoint on the average percentage dry bacteria at Treatment within each treatment (average was calculated among the technical replicates of each biological replicate). **Treatment** (The abiotic treatment), **Term** (predictor used and residuals), **Df** (Degrees of freedom), **Sum Sq** (sum of squares), **F value** (F stats), and **Pr(>F)** (P-value)

(C)

| Treatment | Comparison | diff | P |
|---|---|---|---|
| 32 °C | 7 days - 0 | 9.487 | 0.100 |
| 32 °C | 14 days - 0 | 9.380 | *0.084* |
| 32 °C | 21 days - 0 | 18.179 | **< 0.001** |
| 32 °C | 42 days - 0 | 16.896 | **0.002** |
| 22 °C | 7 days - 0 | -0.456 | 1.000 |
| 22 °C | 14 days - 0 | 11.396 | *0.061* |
| 22 °C | 21 days - 0 | 11.321 | *0.063* |
| 22 °C | 42 days - 0 | 14.856 | **0.012** |
| Drought | 7 days - 0 | 9.749 | 0.707 |
| Drought | 14 days - 0 | 15.492 | 0.343 |
| Drought | 21 days - 0 | 5.798 | 0.941 |
| Drought | 42 days - 0 | 9.461 | 0.727 |

**Notes**: Post-hoc Dunnett test to compare later timepoints with timepoint “0”. **Treatment** (abiotic treatment), **Comparison** (It compares timepoint 0 against the later timepoints), **Diff** (The mean differences between the percentage of dry bacteria in those timepoints), **P** (P-value)

**Table S8**: Statistical models to assess the effect of Treatment and timepoint in the percentage of dry bacteria in mixtures in survival experiments (soil) using (A) Two-Way ANOVA, (B) One-Way ANOVA, (C) Post-Hoc Dunnett test.

(A)

| Term | Df | Sum Sq | F value | Pr(>F) |
|---|---|---|---|---|
| Treatment | 3 | 603.687 | 2.496 | *0.068* |
| Timepoint | 3 | 1426.150 | 5.898 | **0.001** |
| Treatment:Timepoint | 9 | 1825.807 | 2.517 | **0.016** |
| Residuals | 62 | 4997.581 | NA | |

**Notes:** Linear model followed by a Two-Way ANOVA to evaluate the effect of the Treatment and the Timepoint on the average of the percentage of dry (the average was determined among technical replicates of each biological replicate or soil tube). **Term** (predictor used and residuals), **Df** (Degrees of freedom), **Sum Sq** (sum of squares), **F value** (F stats), and **Pr(>F)** (P-value)

(B)

| Treatment | Term | Df | Sum Sq | F value | Pr(>F) |
|---|---|---|---|---|---|
| 22 °C | Timepoint | 3 | 555.635 | 7.278 | **0.003** |
| 22 °C | Residuals | 16 | 407.153 | NA | |
| 32 °C | Timepoint | 3 | 1084.489 | 12.231 | **< 0.001** |
| 32 °C | Residuals | 15 | 443.326 | NA | |
| Drought | Timepoint | 3 | 132.420 | 0.170 | 0.915 |
| Drought | Residuals | 15 | 3886.764 | NA | |
| 4 °C | Timepoint | 3 | 1479.414 | 30.308 | **< 0.001** |
| 4 °C | Residuals | 16 | 260.338 | NA | |

Notes: **Treatment** (The abiotic treatment), **Term** (predictor used and residuals), **Df** (Degrees of freedom), **Sum Sq** (sum of squares), **F value** (F stats), and **Pr(>F)** (P-value)

(C)

| Treatment | Comparison | diff | P |
|---|---|---|---|
| 32 °C | 3 months - 0 | 7.831 | *0.094* |
| 32 °C | 6 months - 0 | 12.861 | **0.005** |
| 32 °C | 12 months - 0 | 21.403 | **< 0.001** |
| 22 °C | 3 months - 0 | 0.701 | 0.992 |
| 22 °C | 6 months - 0 | 4.916 | 0.316 |
| 22 °C | 12 months - 0 | 13.244 | **0.002** |
| 4 °C | 3 months - 0 | 17.156 | **< 0.001** |
| 4 °C | 6 months - 0 | -6.114 | *0.073* |
| 4 °C | 12 months - 0 | 6.212 | *0.068* |
| Drought | 3 months - 0 | 5.328 | 0.916 |
| Drought | 6 months - 0 | -0.887 | 1.000 |
| Drought | 12 months - 0 | 4.276 | 0.953 |

**Notes**: **Treatment** (abiotic treatment), **Comparison** (It compares timepoint 0 against the later timepoints), **Diff** (The mean differences between the percentage of dry bacteria in those timepoints), **P** (P-value)

**Table S9:** Mucoid bacteria have higher relative fitness than dry bacteria (A) Linear models assessing the effect of the alfalfa variety and colony morphology on the rhizobia relative fitness, (B) Within-variety effect of morphology. Both account for population structure with genome-wide variation (PC1 and PC2). O: Oneida, S: SW-4107, V: Vernal

(A)

| **Term** | **Sum Sq** | **Df** | **F value** | **Pr(>F)** |
|---|---|---|---|---|
| Morphology | 2.465 | 1 | 8.802 | **0.003** |
| Variety | 0.199 | 2 | 0.356 | 0.701 |
| PC1 | 6.459 | 1 | 23.062 | **< 0.001** |
| PC2 | 9.240 | 1 | 32.993 | **< 0.001** |
| Morphology:Variety | 1.254 | 2 | 2.238 | 0.108 |
| Residuals | 96.066 | 343 | NA | |

Notes**:** We evaluated the effect of Morphology and alfalfa variety after accounting for population structure with PC1 and PC2. We used a linear model with the function lm(), and used the average fitness per strain as the response, followed by ANOVA with type = "II". **Term** (predictor used and residuals), **Sum Sq** (sum of squares), **Mean Sq** (Mean Squares), **F value** (F stats), and **Pr(>F)** (P-value)

(B)

| **Variety** | **Term** | **Sum Sq** | **Df** | **F value** | **Pr(>F)** |
|---|---|---|---|---|---|
| O | Morphology | 0.833 | 1 | 2.381 | 0.126 |
| O | PC1 | 0.681 | 1 | 1.947 | 0.166 |
| O | PC2 | 0.636 | 1 | 1.817 | 0.180 |
| O | Residuals | 39.541 | 113 | NA | |
| S | Morphology | 1.750 | 1 | 6.138 | **0.015** |
| S | PC1 | 0.844 | 1 | 2.960 | *0.088* |
| S | PC2 | 7.602 | 1 | 26.665 | **< 0.001** |
| S | Residuals | 32.215 | 113 | NA | |
| V | Morphology | 0.234 | 1 | 3.008 | *0.086* |
| V | PC1 | 18.565 | 1 | 238.363 | **< 0.001** |
| V | PC2 | 2.926 | 1 | 37.568 | **< 0.001** |
| V | Residuals | 8.801 | 113 | NA | |

**Notes:** Within each alfalfa variety we evaluated the effect of Morphology after accounting for population structure with PC1 and PC2. We used a linear model with the function lm(), and used the average fitness per strain as the response, followed by ANOVA with type = "II". **Term** (predictor used and residuals), **Sum Sq** (sum of squares), **Mean Sq** (Mean Squares), **F value** (F stats), and **Pr(>F)** (P-value)

**Table S10**: Linear models assessing the effect of the alfalfa variety and colony morphology on the (A) shoot biomass, (B) nodule number, and (C) proportion of nodule branching after accounting for population structure with genome-wide variation (PC1 and PC2).

(A)

| **Term** | **Sum Sq** | **Df** | **F value** | **Pr(>F)** |
|---|---|---|---|---|
| Morphology | 0.282 | 1 | 25.302 | **< 0.001** |
| Variety | 0.498 | 2 | 22.344 | **< 0.001** |
| PC1 | 0.321 | 1 | 28.850 | **< 0.001** |
| PC2 | 0.002 | 1 | 0.210 | 0.647 |
| Morphology:Variety | 0.004 | 2 | 0.166 | 0.847 |
| Residuals | 3.820 | 343 | NA | |

Notes**:** We evaluated the effect of Morphology and alfalfa variety after accounting for population structure with PC1 and PC2. We used a linear model with the function lm(), and used the average plant biomass per strain as the response, followed by ANOVA type = "**II**". **Term** (predictor used and residuals), **Sum Sq** (sum of squares), **Df** (Degrees of freedom), **F value** (F stats), and **Pr(>F)** (P-value)

(B)

| **Term** | **Sum Sq** | **Df** | **F value** | **Pr(>F)** |
|---|---|---|---|---|
| Morphology | 1822.286 | 1 | 8.973 | **0.003** |
| Variety | 9068.839 | 2 | 22.327 | **< 0.001** |
| PC1 | 3168.594 | 1 | 15.602 | **< 0.001** |
| PC2 | 763.718 | 1 | 3.760 | *0.053* |
| Morphology:Variety | 1293.377 | 2 | 3.184 | **0.043** |
| Residuals | 69660.565 | 343 | NA | |

Notes**:** as follows: We evaluated the effect of Morphology and alfalfa variety after accounting for population structure with PC1 and PC2. We used a linear model with the function lm(), and used the average nodule number per strain as the response, followed by ANOVA type = "**II**". **Term** (predictor used and residuals), **Sum Sq** (sum of squares), **Df** (Degrees of freedom), **F value** (F stats), and **Pr(>F)** (P-value)

(C)

| **Term** | **Sum Sq** | **Df** | **F value** | **Pr(>F)** |
|---|---|---|---|---|
| Morphology | 0.013 | 1 | 0.885 | 0.347 |
| Variety | 0.035 | 2 | 1.209 | 0.300 |
| PC1 | 0.456 | 1 | 31.448 | **< 0.001** |
| PC2 | 0.055 | 1 | 3.800 | *0.052* |
| Morphology:Variety | 0.002 | 2 | 0.086 | 0.918 |
| Residuals | 4.977 | 343 | NA | |

Notes**:** We evaluated the effect of Morphology and alfalfa variety after accounting for population structure with PC1 and PC2. We used a linear model with the function lm(), and used the average proportion of branched nodules per strain as the response, followed by ANOVA type = "**II**". **Term** (predictor used and residuals), **Sum Sq** (sum of squares), **Df** (Degrees of freedom), **F value** (F stats), and **Pr(>F)** (P-value)

**Figure S1:** Principal component analysis using the SNPs present in all the strains. (A) Screeplot showing the 10 principal components that explain the highest amount of variation. (B) A biplot showing the first two components: the blue dots are the mucoid strains, and the yellow dots are the dry strains. See more information in Table S1

(A)

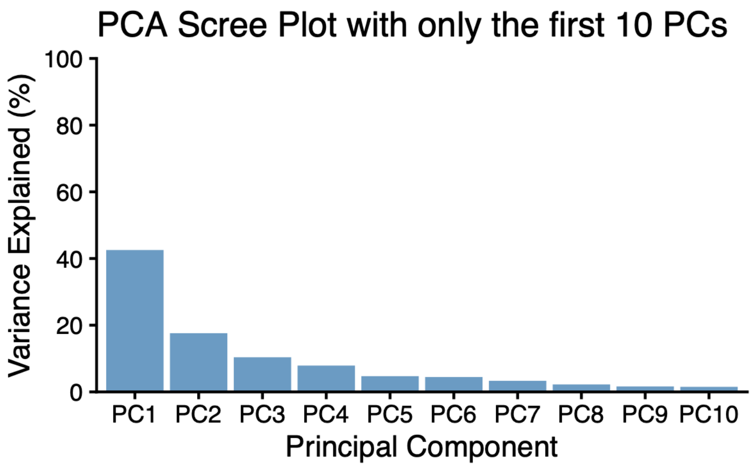


(B)

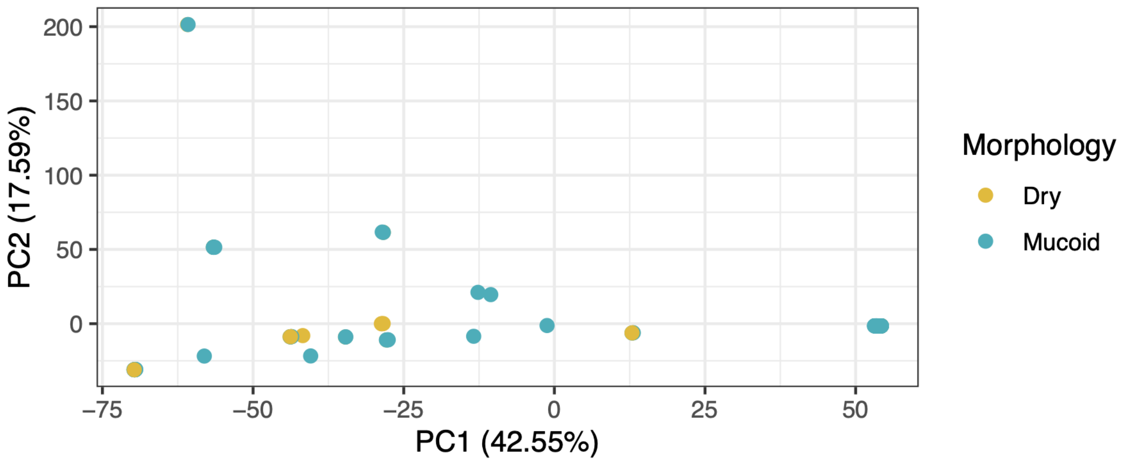

**Figure S2:** Characterizations of the morphologies inside individual nodules collected from plants inoculated with dry and mucoid bacteria. We co-inoculated 4 pairs of dry and mucoid strains in alfalfa plants (7 reps each). Then we sampled, crushed, diluted, and plated 70 nodules per pair, for a total of 280 nodules, to determine whether nodules had dry-only, mucoid-only, or mixed morphologies as seen in (A). In Figure (B), the x-axis shows the strains from the four pairs, and the y-axis shows the proportion of nodules in each category. We have removed the nodules that harbored non-rhizobia-like morphologies or from which nothing grew. For numeric values, see Table S2

(A)

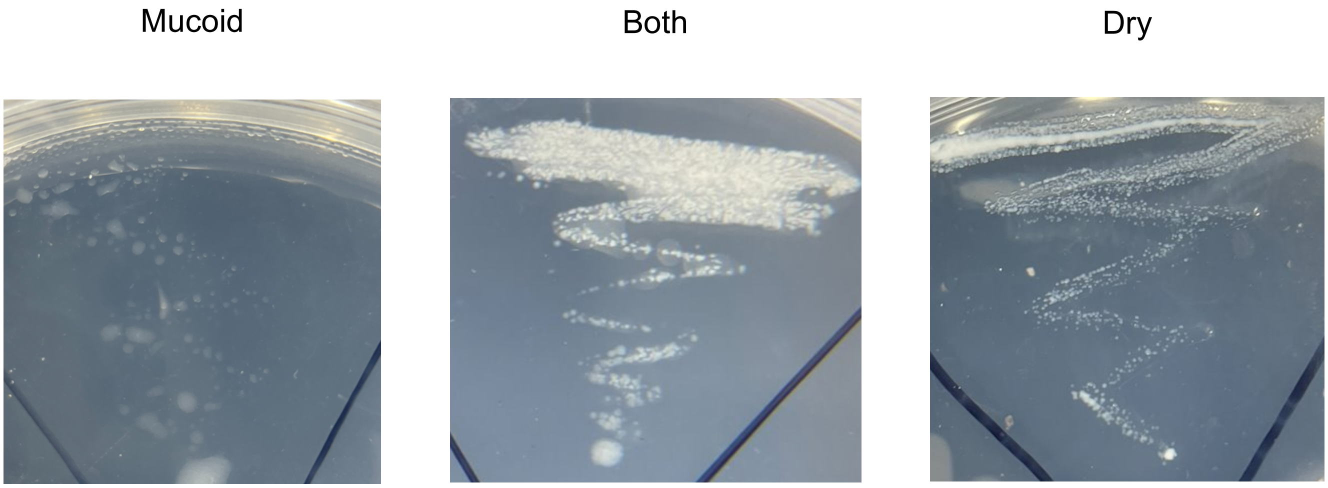


(B)

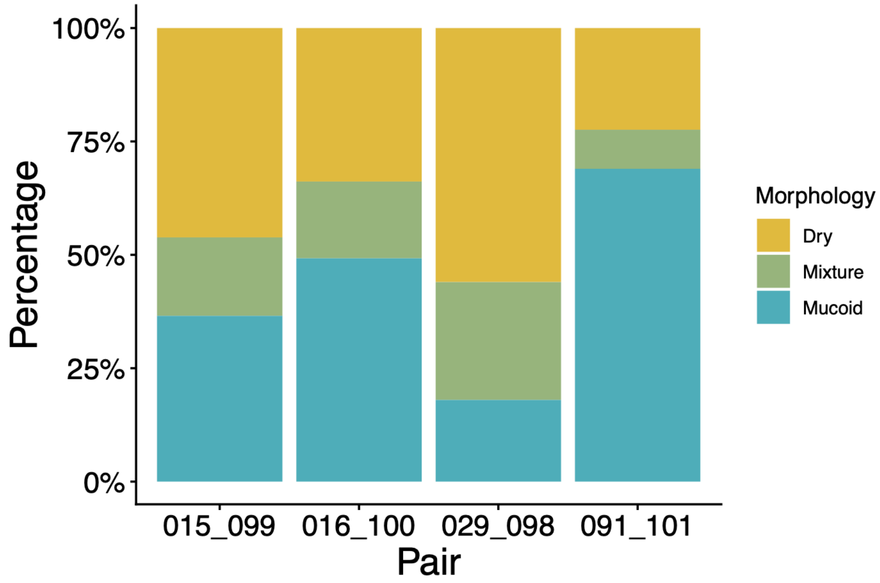

**Figure S3:** Mucoid rhizobia have higher Normalized glucose equivalent (Glucose/$OD_{600}$) when comparing (A) the co-isolated strains and (B) the whole collection. Pairs refer to the two morphologies co-isolated together from an individual well from the several 96-well plates sent by the GALT Prospector Company. More information about the co-isolated strains is in Materials and Methods and Dataset S1. Each filled circle in (B) represents one strain, and there are 115 of them (strain 121 did not grow, and strain 116 was not phenotyped). To see whether morphology affects the normalized glucose equivalent, refer to Table S3.

(A)

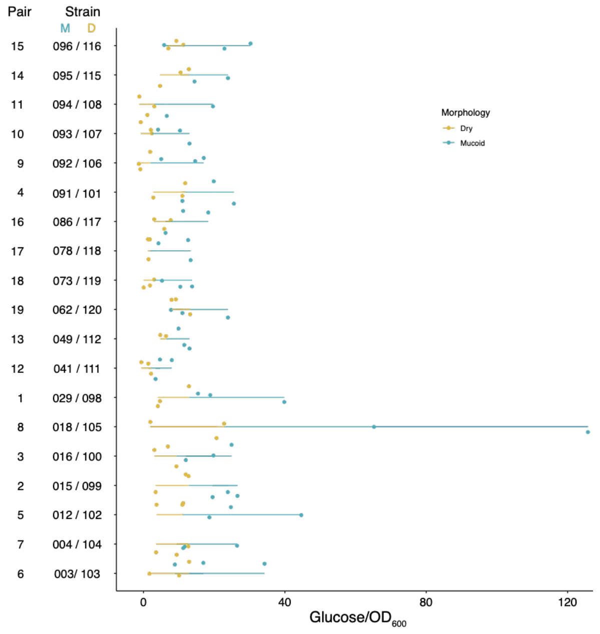


(B)

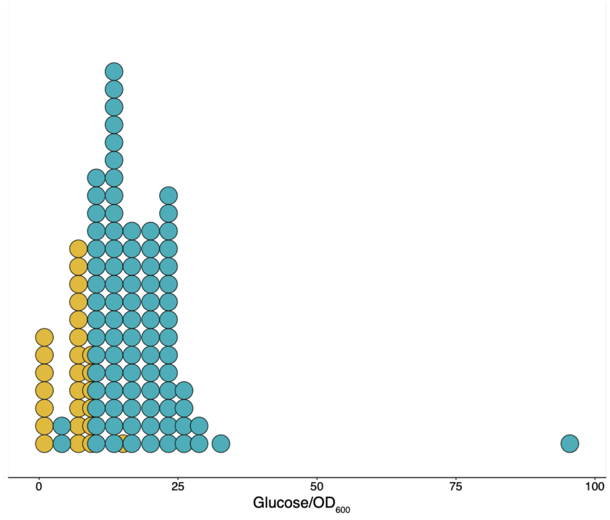

**Figure S4:** Dry and Mucoid populations under the same abiotic treatment show similar trends in both the growth and survival experiments. We grew 27 dry strains to form the dry population and 27 mucoid strains to form the mucoid population. These two populations were separately inoculated into soil tubes, exposed to 4 environmental conditions, harvested, and the population size was measured at 4 timepoints for growth (7, 14, 21, and 42 days) and 3 timepoints for survival (3, 6, and 12 months). We have log10-transformed the population size, which now represents the y-axis. For the “growth” experiment, the rhizobial population did not grow at 4°C even after 7 months, which was the last timepoint assessed for that treatment.

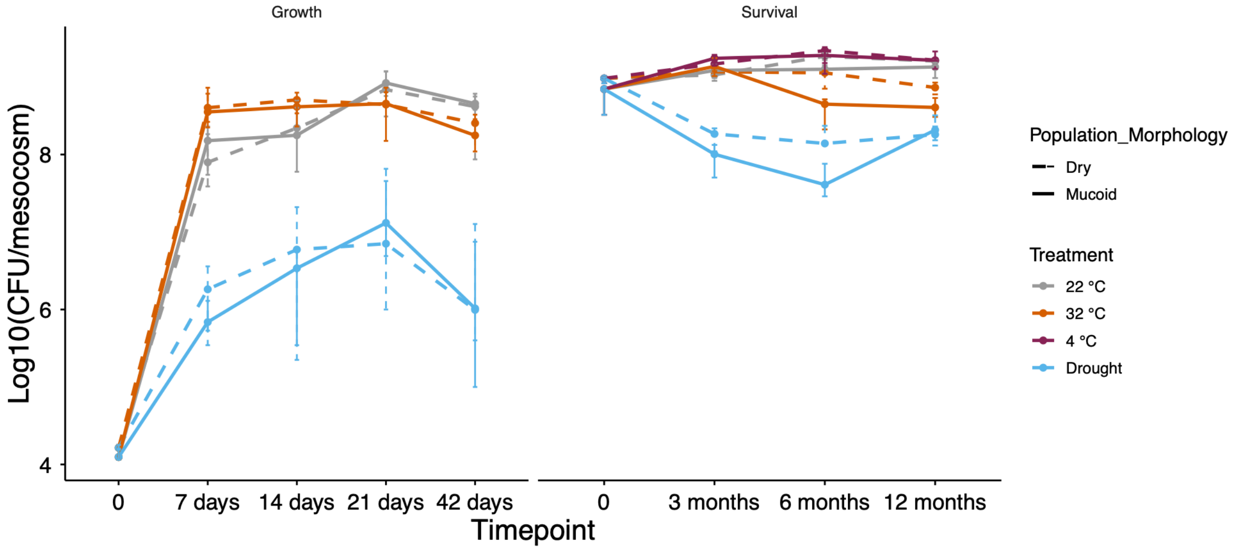

**Figure S5:** Both the survival and growth experiments in soil using the mucoid population show that dry bacteria sporadically show up in soil. (A) Population size in the growth treatment, (B) Population size in the survival treatment. (C) Percentage of dry bacteria in the Growth treatment, (D) Percentage of dry bacteria in the Survival treatment

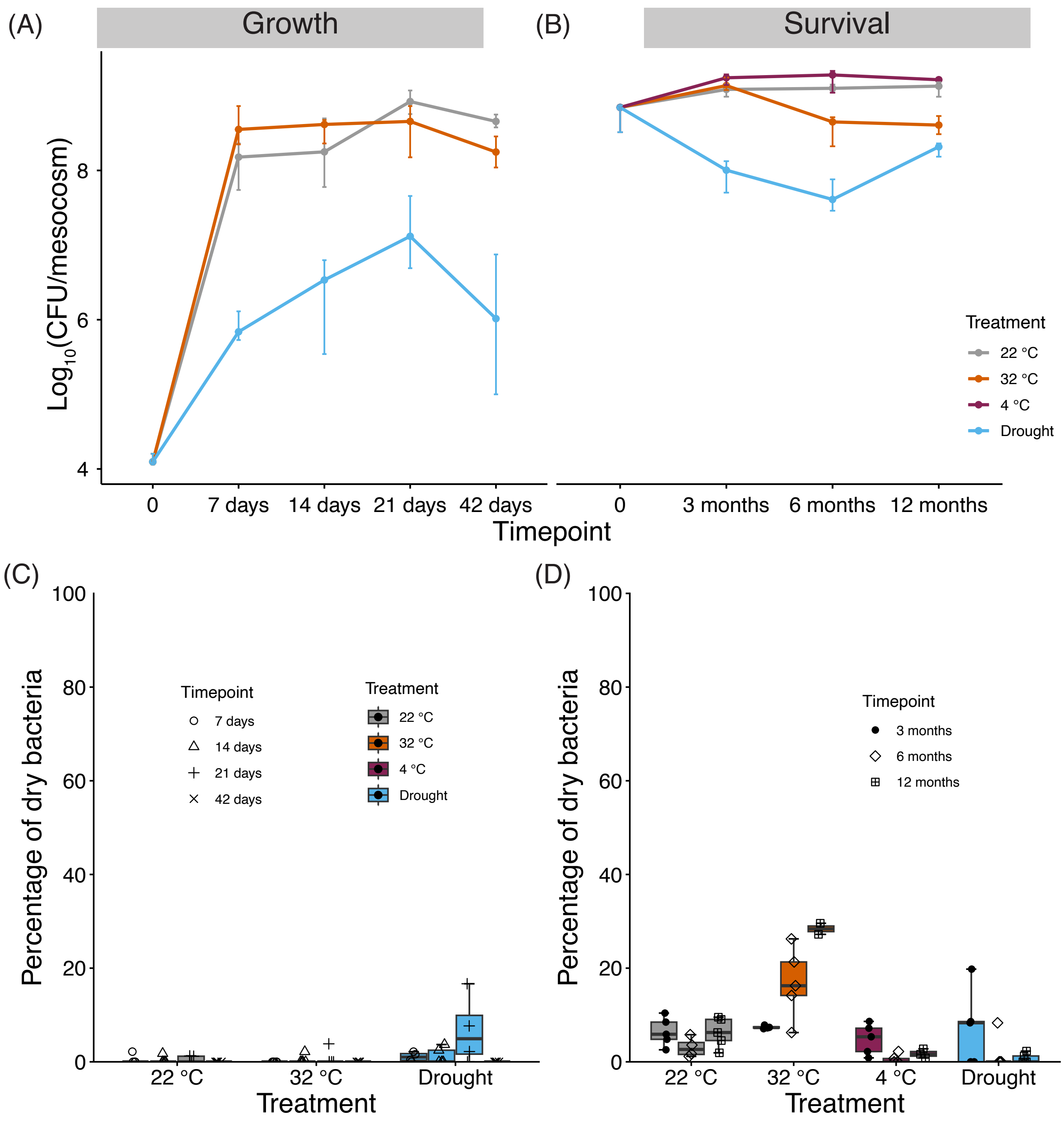

**Figure S6:** In dual inoculation experiments with pairs of dry and mucoid bacteria in the alfalfa variety Oneida, it was found that there is no consistency in whether dry or mucoid bacteria are at higher frequencies in nodule pools. The red dashed line represents the initial percentage of dry bacteria in the inoculum.

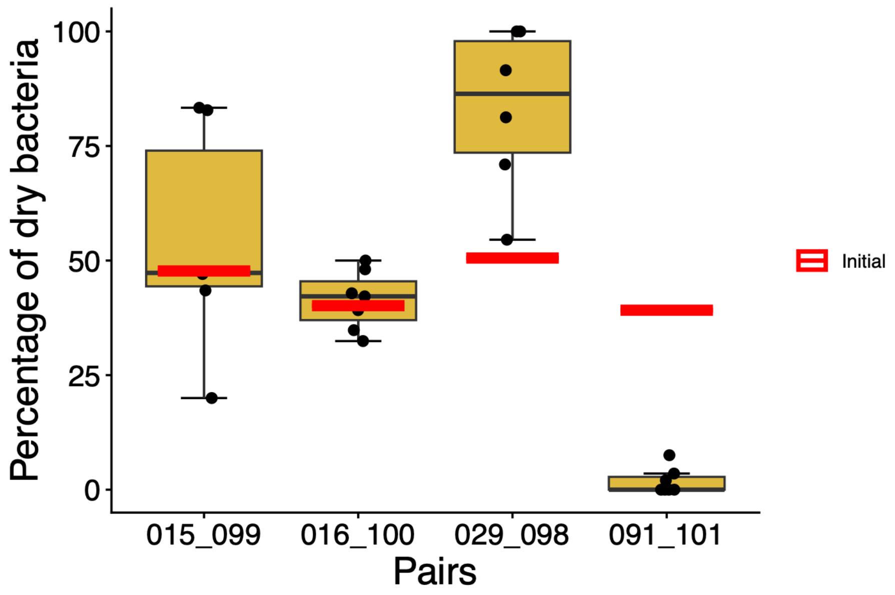

**Figure S7**: Sporadically, dry bacteria would be recovered from nodules of plants inoculated uniquely with a mucoid strain. Red dots represent the plants that were inoculated with mucoid bacteria, and the y-axis represents the percentage of dry bacteria found in those plants.

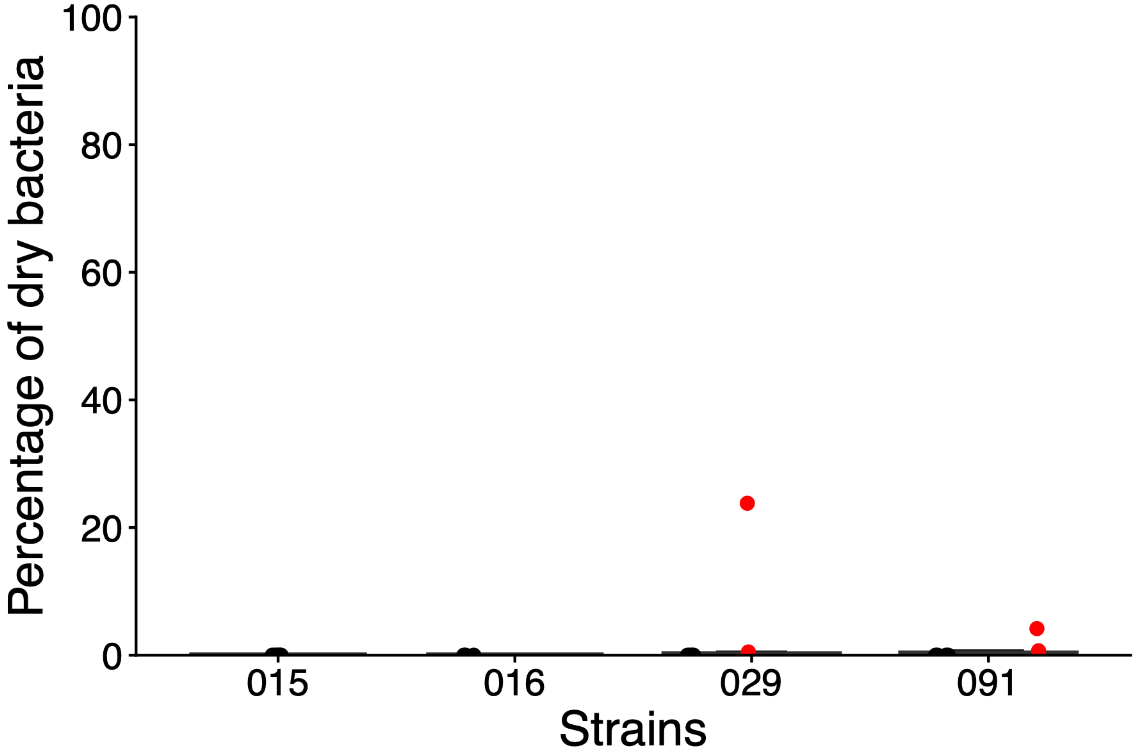

**Figures S8:** Morphology does not have a significant effect on the proportion of branched nodules, and the effect of morphology depends on the variety when evaluating nodule number (A) Nodule number (B) Proportion of branched nodules. Each dot in A and B represents the average of the trait in each strain. The statistics for both traits are in Table S10. Alfalfa varieties Oneida (O), SW-4107 (S), and Vernal (V).

(A)

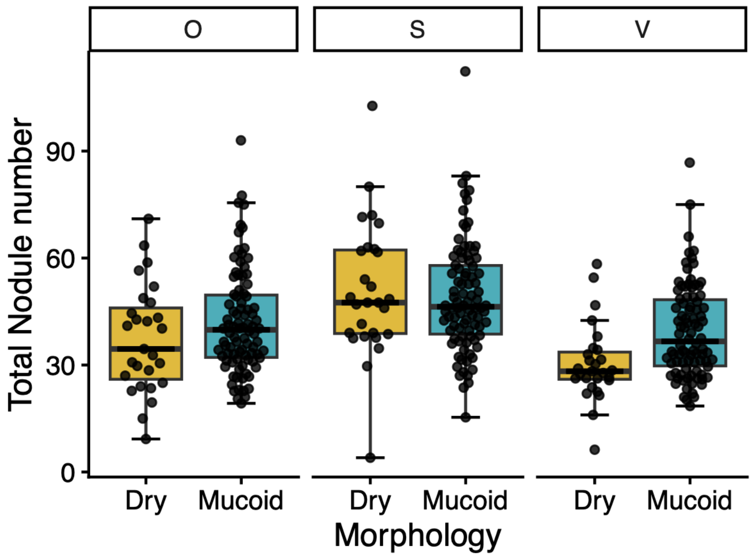


(B)

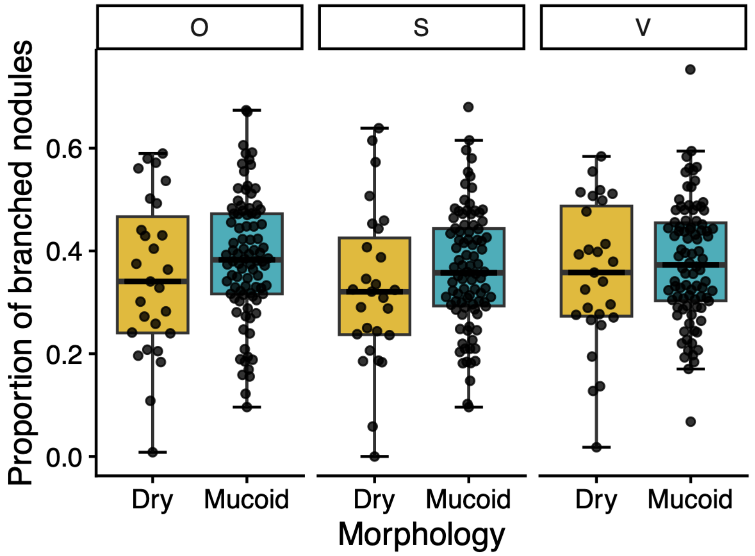

**Dataset S1**: Strains used in this study.

| Strain_Code | Strain | Morphology | Pairs | EPS measurement | Soil mesocosm |
|---|---|---|---|---|---|
| AVT1 | 001 | Mucoid | | Y | - |
| AVT2 | 002 | Mucoid | | Y | M |
| AVT3 | 003 | Mucoid | 6 | Y | M |
| AVT4 | 004 | Mucoid | 7 | Y | M |
| AVT5 | 005 | Mucoid | | Y | - |
| AVT6 | 006 | Mucoid | | Y | - |
| AVT7 | 007 | Mucoid | | Y | - |
| AVT8 | 008 | Mucoid | | Y | - |
| AVT9 | 009 | Mucoid | | Y | - |
| AVT10 | 010 | Mucoid | | Y | - |
| AVT11 | 011 | Mucoid | | Y | - |
| AVT12 | 012 | Mucoid | 5 | Y | M |
| AVT13 | 013 | Mucoid | | Y | - |
| AVT14 | 014 | Mucoid | | Y | - |
| AVT15 | 015 | Mucoid | 2 | Y | M |
| AVT16 | 016 | Mucoid | 3 | Y | M |
| AVT17 | 017 | Mucoid | | Y | - |
| AVT18 | 018 | Mucoid | 8 | Y | M |
| AVT19 | 019 | Mucoid | | Y | - |
| AVT20 | 020 | Mucoid | | Y | - |
| AVT21 | 021 | Mucoid | | Y | - |
| AVT22 | 022 | Mucoid | | Y | - |
| AVT23 | 023 | Mucoid | | Y | - |
| AVT24 | 024 | Mucoid | | Y | - |
| AVT25 | 025 | Mucoid | | Y | - |
| AVT26 | 026 | Mucoid | | Y | - |
| AVT28 | 028 | Mucoid | | Y | - |
| AVT29 | 029 | Mucoid | 1 | Y | M |
| AVT30 | 030 | Mucoid | | Y | - |
| AVT31 | 031 | Mucoid | | Y | - |
| AVT32 | 032 | Mucoid | | Y | M |
| AVT33 | 033 | Mucoid | | Y | - |
| AVT34 | 034 | Mucoid | | Y | - |
| AVT35 | 035 | Mucoid | | Y | - |
| AVT37 | 037 | Mucoid | | Y | M |

| | | | | | |
|---|---|---|---|---|---|
| AVT38 | 038 | Mucoid | | Y | - |
| AVT39 | 039 | Mucoid | | Y | - |
| AVT41 | 041 | Mucoid | 12 | Y | M |
| AVT42 | 042 | Mucoid | | Y | - |
| AVT43 | 043 | Mucoid | | Y | - |
| AVT44 | 044 | Mucoid | | Y | M |
| AVT45 | 045 | Mucoid | | Y | - |
| AVT46 | 046 | Mucoid | | Y | - |
| AVT47 | 047 | Mucoid | | Y | - |
| AVT48 | 048 | Mucoid | | Y | M |
| AVT49 | 049 | Mucoid | 13 | Y | M |
| AVT50 | 050 | Mucoid | | Y | - |
| AVT52 | 052 | Mucoid | | Y | - |
| AVT53 | 053 | Mucoid | | Y | - |
| AVT54 | 054 | Mucoid | | Y | - |
| AVT55 | 055 | Mucoid | | Y | - |
| AVT56 | 056 | Mucoid | | Y | - |
| AVT57 | 057 | Mucoid | | Y | - |
| AVT58 | 058 | Mucoid | | Y | - |
| AVT59 | 059 | Mucoid | | Y | - |
| AVT60 | 060 | Mucoid | | Y | - |
| AVT62 | 062 | Mucoid | 19 | Y | M |
| AVT63 | 063 | Mucoid | | Y | - |
| AVT64 | 064 | Mucoid | | Y | - |
| AVT65 | 065 | Mucoid | | Y | - |
| AVT67 | 067 | Mucoid | | N | - |
| AVT68 | 068 | Mucoid | | Y | - |
| AVT69 | 069 | Mucoid | | Y | - |
| AVT70 | 070 | Mucoid | | Y | - |
| AVT71 | 071 | Mucoid | | Y | - |
| AVT72 | 072 | Mucoid | | Y | - |
| AVT73 | 073 | Mucoid | 18 | Y | M |
| AVT74 | 074 | Mucoid | | Y | - |
| AVT75 | 075 | Mucoid | | Y | - |
| AVT76 | 076 | Mucoid | | Y | - |
| AVT77 | 077 | Mucoid | | Y | - |
| AVT78 | 078 | Mucoid | 17 | Y | M |
| AVT79 | 079 | Mucoid | | Y | - |
| AVT80 | 080 | Mucoid | | Y | - |
| AVT81 | 081 | Mucoid | | Y | - |
| AVT82 | 082 | Mucoid | | Y | - |

| | | | | | |
|---|---|---|---|---|---|
| AVT83 | 083 | Mucoid | | Y | M |
| AVT84 | 084 | Mucoid | | Y | - |
| AVT85 | 085 | Mucoid | | Y | - |
| AVT86 | 086 | Mucoid | 16 | Y | M |
| AVT87 | 087 | Mucoid | | Y | - |
| AVT88 | 088 | Mucoid | | Y | - |
| AVT89 | 089 | Mucoid | | Y | M |
| AVT90 | 090 | Mucoid | | Y | M |
| AVT91 | 091 | Mucoid | 4 | Y | M |
| AVT92 | 092 | Mucoid | 9 | Y | M |
| AVT93 | 093 | Mucoid | 10 | Y | M |
| AVT94 | 094 | Mucoid | 11 | Y | M |
| AVT95 | 095 | Mucoid | 14 | Y | M |
| AVT96 | 096 | Mucoid | 15 | Y | M |
| AVT97 | 097 | Dry | | Y | D |
| AVT98 | 098 | Dry | 1 | Y | D |
| AVT99 | 099 | Dry | 2 | Y | D |
| AVT100 | 100 | Dry | 3 | Y | D |
| AVT101 | 101 | Dry | 4 | Y | D |
| AVT102 | 102 | Dry | 5 | Y | D |
| AVT103 | 103 | Dry | 6 | Y | D |
| AVT104 | 104 | Dry | 7 | Y | D |
| AVT105 | 105 | Dry | 8 | Y | D |
| AVT106 | 106 | Dry | 9 | Y | D |
| AVT107 | 107 | Dry | 10 | Y | D |
| AVT108 | 108 | Dry | 11 | Y | D |
| AVT109 | 109 | Dry | | Y | D |
| AVT110 | 110 | Dry | | Y | D |
| AVT111 | 111 | Dry | 12 | Y | D |
| AVT112 | 112 | Dry | 13 | Y | D |
| AVT113 | 113 | Dry | | Y | D |
| AVT114 | 114 | Dry | | Y | D |
| AVT115 | 115 | Dry | 14 | Y | D |
| AVT116 | 116 | Dry | 15 | Y | D |
| AVT117 | 117 | Dry | 16 | Y | D |
| AVT118 | 118 | Dry | 17 | Y | D |
| AVT119 | 119 | Dry | 18 | Y | D |
| AVT120 | 120 | Dry | 19 | Y | D |
| AVT121 | 121 | Dry | | Y | D |
| AVT122 | 122 | Dry | | Y | D |
| AVT123 | 123 | Dry | | Y | D |

Notes:
**Strain_Code:** Contains the name of the strains as used in Guha et al., (2026). **Strain**: Contains the version of the name of the strains used in this research paper and in Paillan et al (2025). **Morphology**: The initial morphology assigned based on visual inspection. Mucoid bacteria have a watery-like phenotype, while dry bacteria have a more opaque phenotype. **Pairs**: Strains with the same number belong to the same pair; the numbers were randomly assigned. A pair consists of two strains that were co-isolated together from the same individual well from the GALT-prospector's 96-well plates (See Materials and Methods). **EPS_measurement:** Indicates whether the strain was used for the exopolysaccharide quantification assay. Y (yes), N (No). **Soil mesocosm:** The mucoid strains that were used in the soil mesocosm experiments are listed as M, and the dry strains are listed as D. Only 27 mucoid and 27 dry bacteria out of the 117 strains in the collection were used for this experiment
